# Linking biochemical and cellular efficacy of MERS coronavirus main protease inhibitors

**DOI:** 10.64898/2026.02.20.707097

**Authors:** Van N. T. La, Noa Lahav, Moshe Goldsmith, Mario Rodriguez, Randy Diaz-Tapia, Rebecca Pearl, Briana McGovern, Jared Benjamin, Haim Barr, Kris M. White, Lulu Kang, John D. Chodera, David D. L. Minh

## Abstract

Compounds that bind to the Middle East Respiratory Syndrome Coronavirus (MERS-CoV) main protease (MPro) often produce biphasic concentration-response curves (CRCs) in biochemical assays; low concentrations activate the enzyme and high concentrations inhibit it. This biphasic behavior complicates data analysis. Here, we compare three approaches to data analysis: fitting the Hill equation to the activation phase, fitting it to the inhibition phase, and fitting an enzyme kinetics model that incorporates dimerization and ligand binding to the complete CRC. In the latter case, cellular efficacy is predicted by extrapolating the model to high enzyme concentrations. For compounds in our drug lead series, all three procedures yield inhibitory concentrations that are correlated with live-virus antiviral assays. The latter procedure provides the most accurate forecast of cellular efficacy rank. These data analysis procedures may be valuable for antiviral drug discovery against MERS-CoV MPro and other enzymes with similar kinetics.

## 1 Introduction

The Middle East Respiratory Syndrome Coronavirus (MERS-CoV) is a serious threat to global health. The virus was first identified in Saudi Arabia in 2012^1^ and has caused sporadic outbreaks, predominantly in the Middle East, Africa, and South Asia. According to the WHO, no vaccine or antiviral treatment has been approved for MERS-CoV.^2^ The virus has evolved between 2015 and 2019^3^ and further evolution could produce increased transmissibility. Given this possibility and the alarmingly high fatality rate of 35%,^4^ MERS-CoV could lead to large-scale mortality.

Many drug discovery efforts for coronaviruses have focused on identifying compounds that inhibit the main protease (MPro). ^5–11^ MPro is essential to the life cycle of coronaviruses. It is one of 16 non-structural proteins produced upon viral entry into host cells, forming part of the replicase-transcriptase complex responsible for genomic RNA replication and subgenomic mRNA synthesis.^12,13^ Inhibiting MPro disrupts the viral replication cycle,^5,9,14^ facilitating its clearance by the immune system.

In drug discovery campaigns focusing on enzyme inhibitors, concentration response curves (CRCs) that measure the progress of a catalyzed reaction as a function of inhibitor concentration can be a key part of the assay cascade. Improving potency of enzymatic inhibition is one of the most direct objectives of structure-based drug design. While it is possible to forego an enzyme inhibition assay and direct test inhibitors in a cell-based antiviral activity assay, the former generally has fewer safety risks, is less expensive, and is less subject to biological variability. Moreover, cell-based assays can introduce confounding factors, such as membrane permeability and active efflux pumps, that can confuse structure-activity relationships.

Unfortunately, MERS-CoV MPro inhibition assays often show biphasic behavior that complicates their interpretation. MPro is most active as a dimer,^15^ but analytical ultracentrifugation shows that its dissociation constant (*K*_*d*_) is 52 *µ*M,^16^ weaker than MPro from SARS-CoV (6 *µ*M)^17^ and SARS-CoV-2 (7 *µ*M).^18^ Due to the high fraction of enzyme in the relatively inactive monomeric form, ligand-induced dimerization ^16,19^ produces biphasic CRCs, also known as activation-inhibition CRCs, in biochemical assays performed at low enzyme concentrations.^16^ Ligand binding to one monomer can trigger dimerization, locking the catalytic site in an active conformation that stabilizes hydrogen bonding across the dimer interface to the N-terminal serine of the opposite subunit (c.f. Fig. 6 of Nguyen et al. ^20^). If ligand concentrations are low, the other monomer is usually available to bind to substrate and produce product, leading to an overall increase in the catalytic rate. At high ligand concentrations, both monomers are occupied by ligand and enzyme catalysis decreases. For such biphasic curves, the traditional four-parameter Hill equation - which includes bottom response, top response, IC50, and Hill slope - does not fit the complete curve. Thus, it has been unclear how to fit models to these data and how to interpret model parameters for the evaluation of antiviral compounds targeting MERS-CoV MPro and other enzymes that produce biphasic CRCs. Biphasic CRCs have been reported for a noncovalent inhibitor of MERS-CoV MPro^16^ and a reversible covalent inhibitor of mutant SARS-CoV-2 MPro with a weakened dimerization affinity.^21^

Here, we evaluate three possible procedures to interpreting these biphasic CRCs. One is to ignore the activation phase and fit the Hill equation to the inhibition phase. This yields what we will refer to as the *inhibition pIC50*. In another, the inhibition phase is also extracted, but instead of fitting four parameters, the top response is set by the negative control (no inhibitor) and the three remaining parameters are estimated. This procedure essentially assumes that there is no ligand-induced dimerization. We refer to the pIC50 obtained from this fit as the *control pIC50*. A third procedure is based on fitting an enzyme kinetics model that incorporates both dimerization and ligand binding that we recently introduced. ^22^ This model can produce biphasic CRCs that fit to the entire curve without ignoring any data. As it does not explicitly incorporate time dependence, the model is applicable to noncovalent and reversibly covalent inhibitors. Here, we develop a protocol for fitting the model to a large number of CRCs. After fitting the model, we predict CRCs at high enzyme concentrations (reflecting cellular conditions), yielding the *dimer pIC50*. The three procedures are evaluated based on the correlation between different pIC50s and pEC50s in a live-virus antiviral assay.

## 2 Methods

### 2.1 Assays

CRCs were measured in both biochemical enzymatic activity and live-virus antiviral assays, as reported in the AI-driven Structure-enabled Antiviral Platform (ASAP) Discovery Consortium protocols.io repository of experimental protocols.^23^

#### 2.1.1 Biochemical enzyme activity

Biochemical CRCs were obtained by the MERS-CoV MPro fluorescence dose response for antiviral testing protocol^24^ and variants with different concentrations of enzyme, substrate, and inhibitor. The protocol is similar to that described for SARS-CoV-2 MPro, ^25^ but applied to MERS-CoV MPro. Two categories of experiments were performed. In the first category, the enzyme concentration was fixed and the response was measured as a function of substrate concentration. Datasets from this category are referred to as enzyme-substrate (ES) datasets, comprising three datasets with enzyme concentrations at 25, 50, and 100 nM, respectively. Substrate concentrations were 50, 150, 350, 550, 750, 950, 1150, and 1350 nM. For ES datasets, six replicates were measured at each substrate concentration. In the second category, both enzyme and substrate concentrations were fixed while inhibitor concentrations were varied. Datasets from this category are referred to as enzyme-substrate-inhibitor (ESI) datasets. For thirteen inhibitors, CRCs were measured under four conditions (ESI4c): 50 nM enzyme, 150 nM substrate; 100 nM enzyme, 50 nM substrate; 100 nM enzyme, 750 nM substrate; and 100 nM enzyme, 1350 nM substrate. Inhibitor concentrations were 50, 100, 194, 388, 776, 1552, 2488, 7463, 12440, 24880, 49750, and 99500 nM. For 85 inhibitors, CRCs were measured with 50 nM of enzyme and 550 nM of substrate (ESI1c), while the inhibitor concentrations were 0.888, 2, 4, 15, 50, 133, 460, 1227, 2488, 9950, 32340, and 99500 nM. In the ESI datasets, two replicates were measured at each inhibitor concentration. The ESI1c data were obtained by the reported protocol^24^ and ES and ESI4c experiments were performed analogously, but with different concentrations and with fluorescence measurements at different times. Fluorescence was measured every 2 minutes for 2 hours for the ES and ESI4c datasets and once after 60 minutes in the ESI1c dataset. Normalized data are provided in Tables ES, ESI4c, and ESI1c of the Supplementary Information.

Inhibitors were part of a drug discovery campaign for MPro inhibitors targeting both MERS CoV and SARS-CoV-2 conducted by ASAP. Compounds were synthesized by Enamine (Ukraine). The majority of compounds were designed as noncovalent inhibitors, but some have nitrile groups that could form reversible covalent bonds. Complete CRCs and ASAP identifiers are available in Figure S6 and an Excel spreadsheet in the Supplementary Information. Chemical structures have been deposited to ChEMBL 37.

#### 2.1.2 Live-virus antiviral assay

The Live-virus MERS-CoV Vero-TMPRSS2 with PgP Inhibitor Antiviral Screening Assay^26^ was performed at the Icahn School of Medicine at Mount Sinai. All assays were performed at Biosafety Level 3 (BSL-3) in the Emerging Pathogens Facility (EPF).

Vero-TMPRSS2 cells were seeded in 96-well plates at 2,000 cells per well in 10% growth media supplemented with puromycin the day before the assay and incubated at 37 °C and 5% CO_2_. Two hours before infection, cells were treated with 100 *µ*L of a 1 to 3 dilution series of antiviral hits in 2% infection media supplemented with PgP-inhibitor. Dilutions were performed using a Tecan D300e (Tecan). Concentrations of antiviral hits were 50% higher than the target concentrations to account for infection volume. DMSO and uninfected controls were also included on each plate.

Plates were then transferred to the BSL-3 and appropriate wells were infected with MERS-CoV/EMC/2012 at MOI 0.5 in 50 *µ*L of 2% infection media supplemented with PgP-inhibitor, bringing the dilution series to the target concentrations. Plates were then incubated for 48 hours at 37C 5% CO_2_.

48 hours post infection, supernatants were removed from the wells and replaced with 100ul of 4% formalin and incubated for 15 minutes. Outer surfaces of the plates were decontaminated; plates were double bagged, removed from the facility and left to fumigate for 48 hours. Plates were then immunostained using MERS-CoV nucleoprotein (NP) antibody (SinoBiological #40068-RP01) with a DAPI counterstain (Total Cells). Plates were analyzed using a Cytation1 (Biotec). Infectivity was measured by the accumulation of viral N protein (Infected Cells; 488nm). Percent infection was quantified as ((Infected Cells/Total Cells) - Background) * 100, with DMSO control readouts as 100% infection. Data was fit using nonlinear regression and IC50s for each experiment were determined using GraphPad Prism v10.0.0 (San Diego, CA).

All data are included in the Supplementary Information.

#### 2.1.3 Mass photometry

Mass photometry measurements were acquired using a OneMP mass photometer (Refeyn Ltd, Oxford, UK). Microscope coverslips (No. 1.5, 24 × 50 cat 0107222, Marienfeld) were cleaned by sequential sonication in 50% isopropanol (HPLC grade)/Milli-Q H2O and Milli-Q H2O (5 min each), followed by drying with a clean nitrogen stream. Gaskets (MP-CON-21022, Refeyn) were placed at the center of the coverslip, where each well was used for one measurement. Fresh buffer (16 *µ*L) filtered using a 10 kDa Amicon centrifugal filter (Millipore) was first loaded into the coverslip well and used to identify and secure the focal position for the measurement using the “droplet-dilution” mode. For each acquisition, 4 *µ*L of diluted protein solution was mixed with the loaded buffer several times, and data was collected for 2 min, at RT, using a regular window size. Monomer and dimer peaks of BSA (Sigma) and a purified protein BOS (56kDa) were used for calibration. Data acquisition was performed using AcquireMP (Refeyn 2025 R1.2) and data analysis was performed using DiscoverMP (Refeyn 2025, vR1).

### 2.2 Bayesian regression for biochemical enzyme activity

We fit our enzyme kinetics model (Figure 1) that includes dimerization and inhibition^22^ to the ES and ESI datasets. In comparison to the quadratic fits that demonstrate that coronavirus MPro is an obligate homodimer,^21,27^ our model accounts for a more comprehensive set of states and catalytic rates. A complete set of equations and descriptions of numerical solutions to the equations are included in our previous publication.

**Figure 1.**
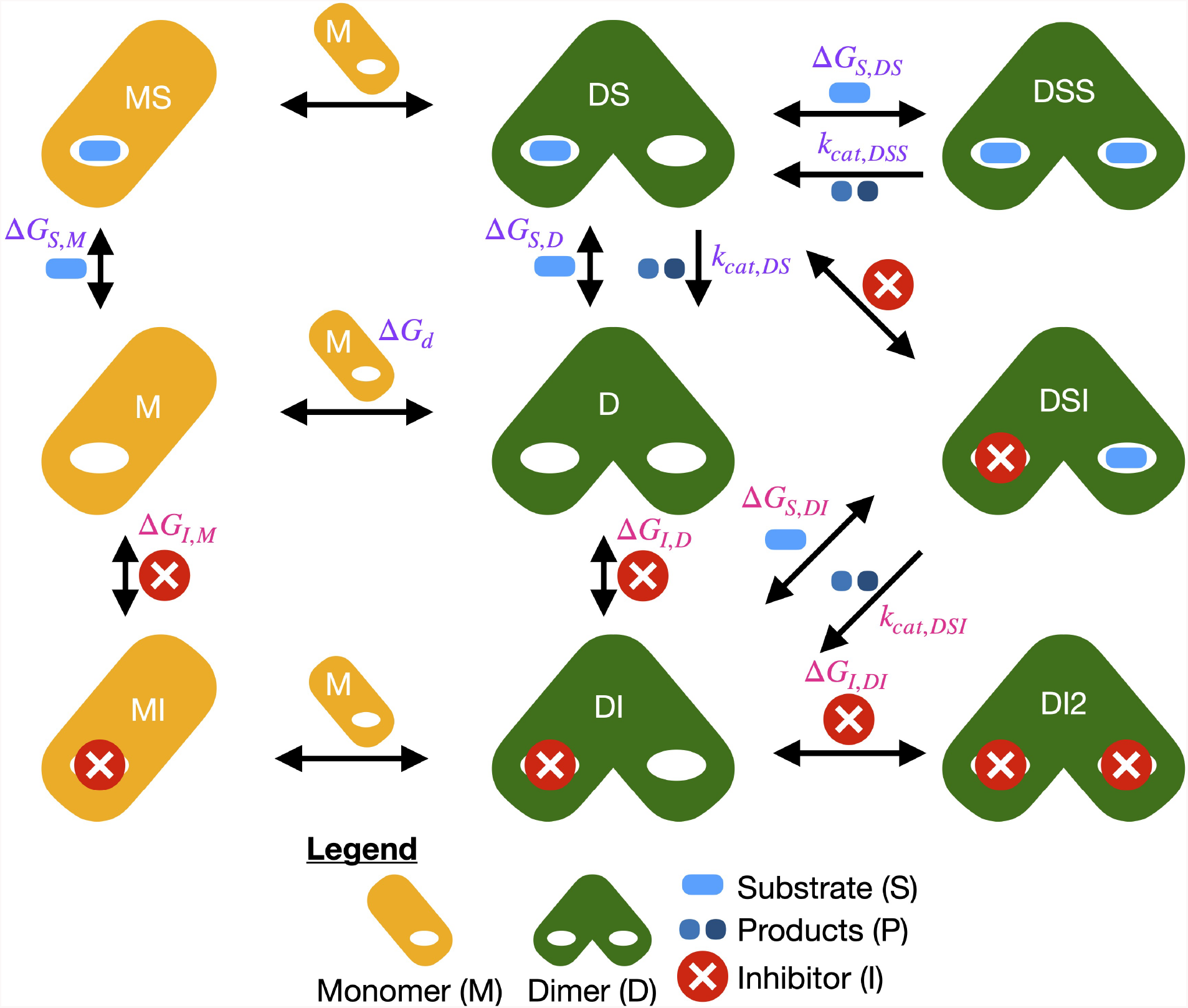
Enzyme kinetics model with dimerization and competitive inhibition. Protein is shown as an orange rounded rectangle for the monomer (M) or pair of overlapping green rounded rectangles for the dimer (D). Species on top of arrows are added going right / removed going left. Species to the right of arrows are added going down / removed going up. Free energies are forward for the direction that leads to more complex species. Rate constants (*k*_*c*_*at*) depend on the dimerization and ligand binding. Parameters shared between different inhibitors are colored grape and inhibitor-specific parameters colored salmon.

#### 2.2.1 Parameters

The objective of our Bayesian regression was to infer the following parameters,

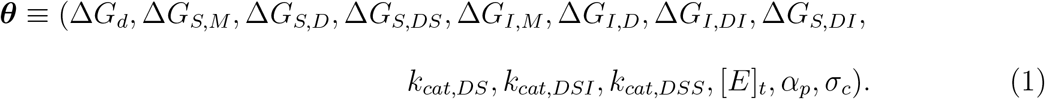

The enzyme kinetics model, which is based on a rapid equilibrium assumption, has thermodynamic (Δ*G*) and kinetic (*k*_*cat*_) parameters as previously described. ^22^ Δ*G* are the binding free energies of species including the MPro monomer (M), MPro dimer (D), substrate (S), inhibitor (I): Δ*G*_*d*_ is a free energy of dimerization; Δ*G*_*S,M*_ is the binding free energy of the substrate to the monomer; Δ*G*_*S,D*_ is the binding free energy of the substrate to the dimer; Δ*G*_*S,DS*_ is the binding free energy of the substrate to the dimer-substrate complex; Δ*G*_*I,M*_ is the binding free energy of the inhibitor to the monomer; Δ*G*_*I,D*_ is the binding free energy of the inhibitor to the dimer; Δ*G*_*I,DI*_ is the binding free energy of the inhibitor to the dimer-inhibitor complex; and Δ*G*_*S,DI*_ is the binding free energy of the substrate to the dimer-inhibitor complex. *k*_*cat*_ are enzyme velocities: *k*_*cat,DS*_ is the velocity of the dimer-substrate complex; and *k*_*cat,DSS*_ is the velocity of the dimer bound to two substrates. The velocity of the monomer-substrate complex is known to be much smaller than the dimer-substrate complex and was therefore assumed to be zero.

Some thermodynamic and kinetics parameters were treated as global, the same for every dataset, and others local, dependent on the inhibitor. The global parameters were the binding free energy of dimerization Δ*G*_*d*_, binding free energies between the enzyme and the substrate Δ*G*_*S*_, and rate constants *k*_*cat,DS*_ and *k*_*cat,DSS*_. The inhibitor-dependent parameters were the binding free energies of the inhibitor binding to the enzyme Δ*G*_*I*_, the binding free energy of the substrate binding to the enzyme-inhibitor complex Δ*G*_*S,DI*_, and the velocity of the enzyme-substrate-inhibitor complex *k*_*cat,DSI*_.

In addition to the thermodynamic and kinetic parameters, our model also uses several local parameters: [*E*]_*t*_, *α*_*p*_, and *σ*_*c*_. [*E*]_*t*_ is the true enzyme concentration. True concentrations may differ from the stated concentrations due to dilution errors or protein degradation. We used one parameter [*E*]_*t*_ for each of the three stated monomer concentrations of 25, 50, and 100 nM; [*E*]_25_ is the true concentration for the solution with a stated concentration of 25 nM, and analogously for 50 and 100. *α*_*p*_ is a scaling factor for all velocities on a given plate *p*. It accounts for differences in velocity calibration due to factors such as plate material or path length variation, instrument lamp intensity or detector sensitivity fluctuations, and sample variations such as pH or buffer evaporation. There was 1 plate for ES, 4 plates for ESI4c, and 45 plates for ESI1c. *σ*_*c*_ is the standard error of the velocity, indexed by *c*, and is assumed to be constant for all points in a CRC. *σ*_*c*_ was also used in our previous work.^22^

#### 2.2.2 Likelihood

For each CRC, the data ***D*** ∈ {*y*_1_, *y*_2_, …, *y*_*n*_} are initial velocities (v, M/min) of the enzymatic reaction. Initial velocities were calculated based on linear regression and normalization to ensure that the same rates are obtained for the same reaction conditions. The total fluorescence response *R* was assumed to the sum of the response of the fluorescent substrate and product. For each species, the fluorescence response is the concentration of the species (*c*_*s*_ for substrate and *c*_*p*_ for product) and its molar response (*r*_*s*_ for substrate and *r*_*p*_ for product),

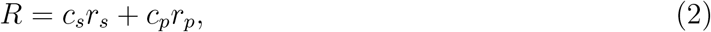

which has the time derivative,

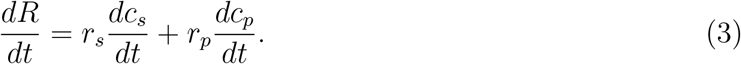

Thus, as substrate is converted into product, the observed slope is *m* = (*r*_*p*_ − *r*_*s*_)*v*, where *v* is the initial velocity of the reaction. For each well, the slope *m* was determined by measuring the response from the biochemical assay every 2 minutes for 10 minutes after addition of substrate and performing ordinary least squares linear regression (linalg.lstsq) as implemented in numpy. *r*_*s*_ was determined by dividing the intercept by the initial substrate concentration. On the master plate with the ES dataset, *r*_*p*_ was calibrated by measuring fluorescence after 21 hours, at which point the substrate was assumed to be completely converted to product. For plates with the ESI4c datasets, *r*_*p*_ for each plate was determined by solving a system of linear equations such that at the same enzyme and substrate concentrations, the initial velocity of the plate and the master plate are equal. Velocities in the ESI1c dataset were normalized to a velocity interpolated from the ES and ESI4c datasets. After fitting the velocities from the ES and ESI4c datasets by Bayesian regression, maximum a posteriori (MAP) parameters were used to estimate a reference velocity *v*_0_, the velocity at 50 nM enzyme and 550 nM substrate; this condition was measured on all of the plates of the ESI1c dataset.

During exploratory data analysis, we identified outliers, many which we attributed to limited solubility at high compound concentrations. Before fitting the model, outliers were removed using a *z*-score test.^28^ A pooled standard deviation was calculated for each CRC in the ESI datasets as,

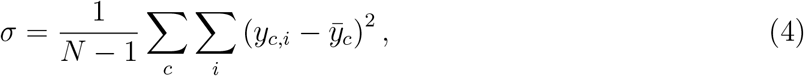

where *y*_*c,i*_ is a measured velocity at a given condition (enzyme, substrate, and inhibitor concentration) and 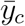 is the sample mean of velocities at the condition. Sums are over the measurements and conditions and *N* is the number of measurements in all conditions (six for ES datasets and two for ESI datasets.) The *z*-score was calculated as,

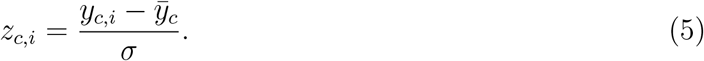

An observation *y*_*c,i*_ is considered an outlier if the absolute value of its corresponding *z*_*c,i*_ score exceeds 2.5. The thresholds of −2.5 and 2.5 correspond to the 2.5th and 97.5th percentiles of the observations in the dataset, respectively. Any outliers present in each CRC were removed before fitting.

Measurements were assumed to follow a normal distribution centered around model-predicted values (scaled by *α*_*p*_), 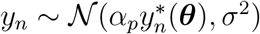. The likelihood of the data ***D*** is given by,

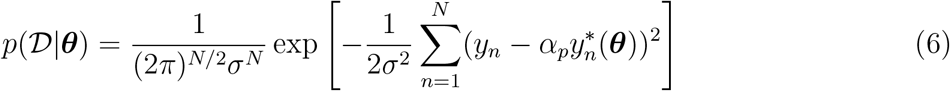

in which the measurement 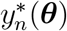 is a function of all parameters in ***θ***, except for *α*_*p*_.

#### 2.2.3 Prior

Assuming that the parameters are independent, the prior *p*(***θ***) is a product of the prior for all parameters, *p*(***θ***) = Π_*i*_ *p*(*θ*_*i*_). Based on the reported value of *K*_*d*_ (52 ± 5 *µ*M)^16^ and the relationship between binding free energy and dissociation constant through Δ*G* = −*RT* ln *K*, the prior of Δ*G*_*d*_ would follow a normal distribution with a mean of −5.9 and a standard deviation of 0.06. However, to reduce the influence of this prior we chose a larger standard deviation,

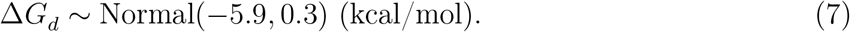

Broad uniform priors were chosen for other binding free energies. The range of Δ*G*_*S*_ was based on *K*_*S*_ between 1 nM and 1 M. The range of Δ*G*_*I*_ was based on *K*_*I*_ between 1 pM and 1 M.

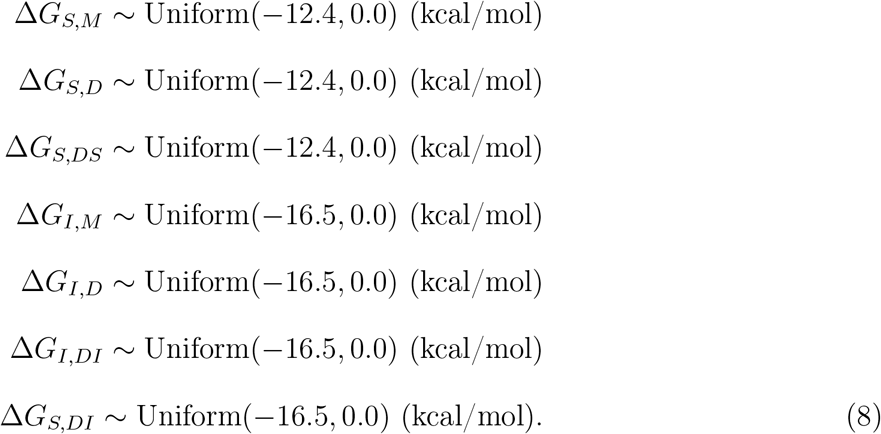

Broad uniform priors were also selected for the kinetic parameters. Based on the reported value of *k*_*cat*_ (0.2 ± 0.02 *min*^−1^),^16^ we chose,

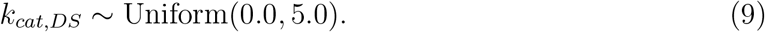

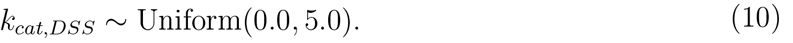

Due to the biphasic behavior observed in CRCs,^16^ we set a higher upper limit for *k*_*cat,DSI*_ ~ Uniform(0.0, 10.0).

For prior of *α*_*p*_, uniform distribution was used,

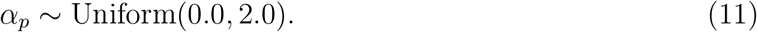

where *p* is the index for the plate. Because the uncertainty in concentration due to sample preparation in biochemical assays has been shown to be approximately 10%,^29^ we used a log-normal prior with 10% uncertainty for the enzyme concentration,

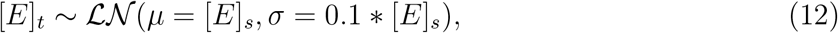

where [*E*]_*s*_ for *s* ∈ {25, 50, 100} nM is the stated value of the enzyme concentration and [*E*]_*t*_ is the true value. The uninformative Jeffreys prior^30^ was used for *σ* of each CRC, as in our previous work.^22^

#### 2.2.4 Sampling from the posterior

As the complexity of the model and large amount of data made global fitting computationally prohibitive with our limited computing resources, we divided the fitting process into several steps, leveraging the posterior distribution from one step to limit the prior of the next.^31^

1. A simplified enzyme kinetics model without inhibitor was fit to the ES dataset to obtain ranges of Δ*G*_*d*_, Δ*G*_*S,M*_, Δ*G*_*S,D*_, Δ*G*_*S,DS*_, *k*_*cat,DS*_, *k*_*cat,DSS*_, and [*E*]_*t*_. As we treated the ES plate as a reference, *α*_*p*_ was set to 1.
2. The full enzyme kinetics model was fit to the ES dataset and curves from *each* inhibitor in the ESI4c dataset. The priors of Δ*G*_*d*_ and Δ*G*_*S*_ were defined based on the minimum and maximum values of these parameters observed in posteriors from step 1.
3. The full enzyme kinetics model was globally fit to the ES dataset and the *full* ESI4c dataset. The priors of Δ*G*_*d*_, Δ*G*_*S*_, and *α*_*p*_ were defined based on the minimum and maximum values of these parameters observed in every posterior from step 2.
4. The full enzyme kinetics model was globally fit to the ES dataset, the *full* ESI4c dataset, and three selected curves from the ESI1c dataset. Priors were defined as in Step 4. The three curves were selected based on criteria described below.
5. When fitting to the ESI1c dataset, global parameters were fixed to the MAP of Step 3 or 4 and local parameters were sampled from the conditional probability.

The three curves in Step 4 were selected based on using the MAP from Step 3 in Step 5. The model from Step 5 did not fit to these three ESI1c curves. Therefore, we incorporated the data in Step 4 in order to obtain a MAP capable of fitting not only ES and ESI4c data, but also the selected ESI1c data.

The No-U-Turn sampler (NUTS) was used to sample from posterior distributions.^32^ NUTS was run for 10,000 samples in steps 1 and 2. Because we observed that the posteriors were already converged by 1,000 samples in steps 1 and 2, we collected 1,000 equilibrated samples in steps 3 and 4. The equilibration time of all the parameters was detected using automated equilibration detection^33^ as implemented in pymbar v4.0.3. ^34,35^

### 2.3 Estimating pIC50s and pIC90s

Inhibitory concentrations were estimated by fitting data with the Hill equation,

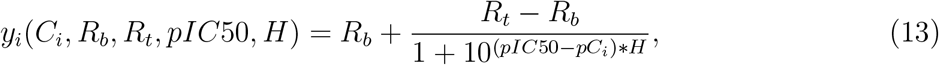

where *R*_*b*_ is the bottom response, *R*_*t*_ is the top response, *pIC*50 is the negative base 10 logarithm of the half maximal inhibitory concentration *IC*50, and *H* is the Hill slope.

As outlined at the end of the introduction, we estimated inhibitory concentrations based on three types of CRCs. For the inhibition pIC50, we first identified the concentration of inhibitor that yields the maximum response. The Hill equation was fit to data at this and higher concentrations. For the control pIC50, the Hill equation was fit to data at this and higher concentrations where the response is less than the negative control (Figure 2).

**Figure 2.**
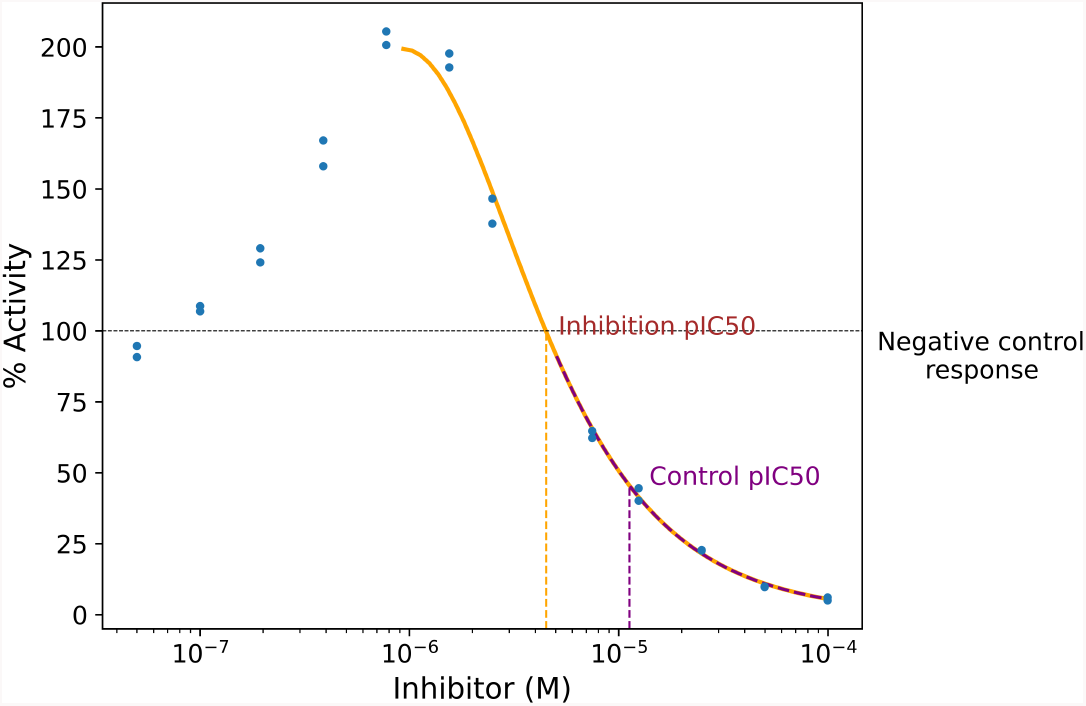
Representative inhibition *IC*50 and control *IC*50.

For the dimer pIC50, we simulated CRCs using the dimer-only kinetic model and fit the Hill equation to the entire curve.^22^ Enzyme and substrate concentrations were drawn from lognormal distributions with a 10% uncertainty. The substrate concentration was selected to be saturating, 1000 times larger than the enzyme concentration. These concentrations were selected by dividing the ESI1c dataset (85 curves) into a training set (45 curves) and a testing set (40 curves). Enzyme concentrations were optimized by minimizing the root mean square deviation between the dimer pIC50s and cellular pEC50s within the training set. This optimization was performed by `scipy.optimize.minimize.`^36^ Optimized concentrations were then applied to estimate dimer pIC50/pIC90 values in the testing set. The reported mean and standard deviation are results from repeating this procedure 100 times.

Velocities were simulated at 50 geometrically distributed inhibitor concentrations between 1 pM to 1 mM and normalized to be between 0 and 100%.

Hill equation parameters were estimated using maximum likelihood estimation. For the biochemical data, estimation was performed using our custom code.^37^ The EC50 is the half maximal effective concentration in cellular assays. Cellular pEC50s for antiviral assays were estimated by fitting the Hill equation to the data using CDD Vault.^38^

Besides IC50, another commonly used metric for assessing the potency of a drug is the *IC*90. This parameter denotes the concentration at which 90% of the enzyme is inhibited. Given the *IC*50, the Hill slope *H*, and a specific percentage F ranging between 0 to 100, representing for the degree of enzymatic inhibition or cellular viability, the *IC*(*F*) can be calculated using the formula 14,^39^

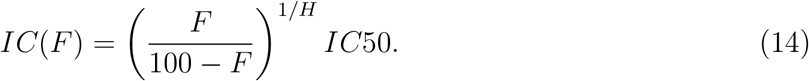

With *F* set to 90, the formula simplifies to,

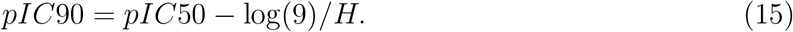

Analogous formulae apply to pEC50 and pEC90 values for cellular assays.

### 2.4 Correlation analysis

Correlation between the pIC50/pIC90 values obtained from different biochemical procedures and cellular pEC50/pEC90 were analyzed by a range of statistical measures, including the Pearson R,^40^ Spearman *ρ*,^41^ Kendall *τ*,^42^ root mean square deviation (RMSD), and adjusted RMSD (aRMSD).^43^ Unlike the Pearson R, which measures linear correlation, the Spearman *ρ* assesses the monotonic relationship between two variables by ranking data points and evaluating how well the ranks correspond, making it robust to nonlinearity. The Kendall *τ* evaluates whether pairs of data points move in the same or opposite directions. RMSD evaluates the absolute differences between predicted and observed values,

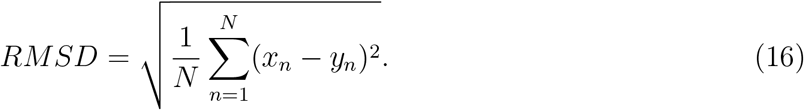

The adjusted RMSD (aRMSD) avoids the effect of systematic bias by normalizing RMSD using the means of the data, providing a location-independent measure of predictive accuracy,

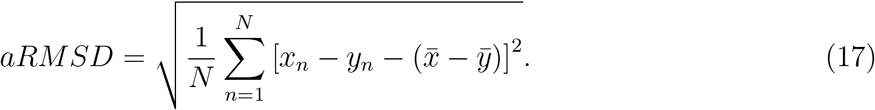

For the dimer pIC50/pIC90, the ESI4c 85 dataset was randomly split into a training set (45 curves) and a test set (40 curves). Enzyme concentrations optimized using the training set were applied to estimate dimer pIC50/pIC90 values in the test set. This procedure was repeated 100 times and the mean and standard deviation of results are reported.

### 2.5 Code

All code is freely available on GitHub.^44^

## 3 Results

Assay linearity and the consistency of the 1-h endpoint readout with initial (0–10 min) velocities were validated using representative time-course traces and plate-wide extrapolation analyses (Appendix A of the Supplementary Information). After performing the described sampling from the Bayesian posterior, Bayesian credible intervals were converged (Appendix B of the Supplementary Information).

### 3.1 Global fitting increases the precision of parameter estimates

Overall, we observed that Bayesian posterior probability distributions became narrower with the inclusion of additional data. In this section, we describe results from Steps 1 through 4. All Step 2 results are with a representative compound, ASAP-0000214.

Marginal distributions of all enzyme-substrate binding free energies are unimodal and nearly independent (Figures 3, S7, and S10). Δ*G*_*d*_ has a relatively small highest density interval (HDI) that is unaffected by the amount of data analyzed. The other parameters are broader in Steps 1 and 2, with Δ*G*_*S,M*_ and Δ*G*_*S,DS*_ approaching the upper limit of weak affinity. All parameters are defined more precisely in Steps 3 and 4. For instance, the 95% HDI of Δ*G*_*S,M*_ in Steps 1 and 2 ranged between −7.5 and 0, whereas in Step 3 and 4, the HDI fell into a narrower range between −5.7 and −4.4. Between Steps 3 and 4, there is no significant difference in posterior distributions of these parameters. In Steps 1 and 2, two-dimensional marginal distributions suggest that pairs of enzyme-substrate binding free energies are nearly independent (Figures S8 and S11). In Steps 3 and 4, however, Δ*G*_*S,M*_ is negatively correlated with Δ*G*_*S,D*_ (Pearson R = −0.75) and positively correlated with Δ*G*_*S,DS*_ (Pearson R = 0.94). Δ*G*_*S,D*_ and Δ*G*_*S,DS*_ are negatively correlated with each other (Pearson R = −0.76) (Figures S17, S18, S20, and S21).

**Figure 3.**
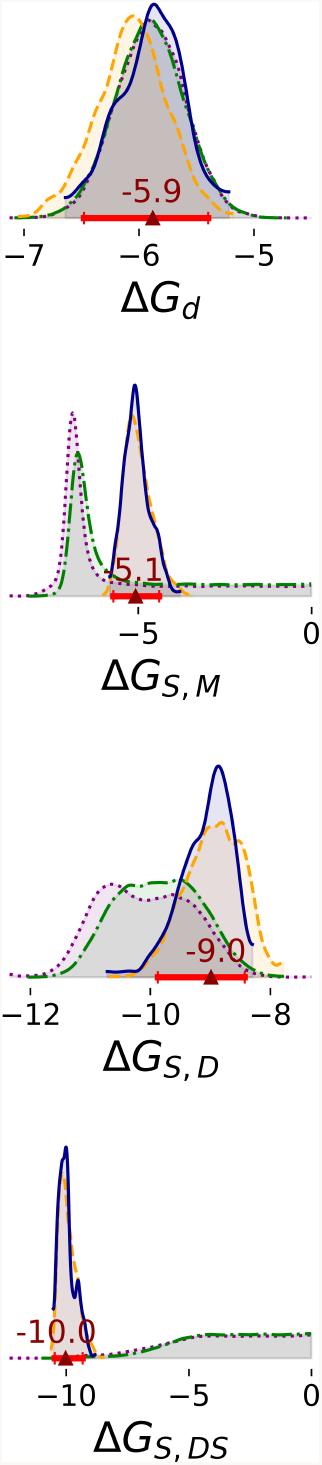
1D marginal distributions of free energies (kcal/mol). Estimates were based on 1,000 MCMC samples generated from the Bayesian posterior in Steps 1 (purple dotted line), 2 (green dashdot line), 3 (orange dashed line), and 4 (blue solid line). Red bars represent 95% HDIs from Step 4. The red triangle marks the median of that posterior.

With additional data, the binding cooperativity of the substrate also becomes more clearly determined (Figure 4). In steps 1 and 2, the estimated difference between Δ*G*_*S,DS*_ and Δ*G*_*S,D*_ is broad (step 1: median 6.5 and 95% HDI [1.7, 11.0] kcal/mol; step 2: median 6.6 and 95% HDI [2.2, 10.0] kcal/mol). The median is positive, suggesting negative cooperativity of substrate binding, with the caveat of low precision. However, with additional data, the posterior is much narrower and the median indicates positive cooperativity of substrate binding (step 3: median −0.99 and 95% HDI [−2.3, 0.57] kcal/mol; step 4 median −0.89; 95% HDI: [−1.9, 0.54] kcal/mol).

**Figure 4.**
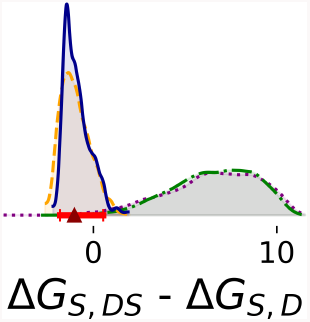
Differences in binding free energies of substrate (kcal/mol). Estimates were based on 1,000 MCMC samples generated from the Bayesian posterior in Steps 1 (purple dotted line), 2 (green dashdot line), 3 (orange dashed line), and 4 (blue solid line). Red bars represent 95% HDIs from Step 4. The red triangle marks the median of that posterior.

Marginal distributions of all enzyme-inhibitor binding free energies are unimodal, and some show significant correlations (Figure S11). In Step 2, Δ*G*_*I,DI*_ reaches the lower bound of strong affinity, indicating that the second binding of the inhibitor to the dimer-inhibitor complex is highly favorable. Correlations can be summarized via Pearson correlation coefficients. In Step 2, the heatmap of the estimated parameters indicates that Δ*G*_*S,DS*_ and Δ*G*_*I,M*_, which have broad posterior distributions, have no correlation with any other parameters (Figure S12). On the other hand, Δ*G*_*I,D*_ displays a strong negative correlation with both Δ*G*_*I,DI*_ and Δ*G*_*S,DI*_, but have no correlation with any other dissociation constants (Figures S12, S13a and S13b). Δ*G*_*S,DI*_ demonstrates a positive linear correlation with Δ*G*_*I,DI*_, although the difference between them is small (Δ*G*_*S,DI*_=Δ*G*_*I,DI*_+1.83, Figure S13c).

Bayesian posterior distributions of most rate constants are broad; data are insufficient to estimate these parameters accurately (Figure 5). For Steps 1 and 2, the marginal of *k*_*cat,DSS*_ has a peak at low rates and a heavy tail that spans the range of the prior. In Steps 3 and 4, the posterior of low rates is significantly decreased. The posterior of *k*_*cat,DSS*_ is flat in Steps 1 and 2, but sharply defined after Steps 3 and 4. For the representative ligand, posteriors of *k*_*cat,DSI*_ are flat despite the inclusion of more data (Figure S14). Additionally, 2D joint marginal distributions show that there is no correlation between rate constants (Figure S15).

**Figure 5.**
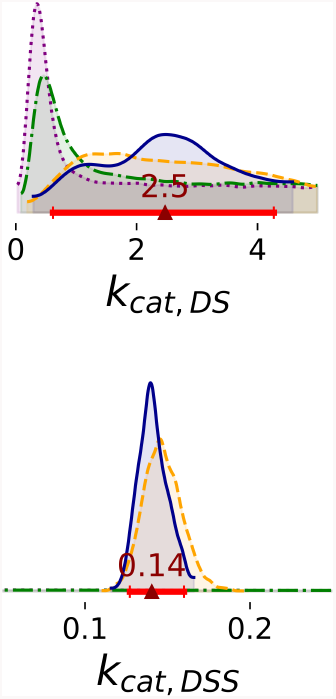
1D marginal distributions of rate constants (min^−1^). Estimates were based on 1,000 MCMC samples generated from the Bayesian posterior in Steps 1 to 4. The posterior distribution of *k*_*cat,DSS*_ is shown zoomed in opposed to the full domain between 0 and 5 observed for Steps 1 and 2. Annotations are analogous to Figure 3.

While individual rates are difficult to determine, the data are sufficient to estimate ratios of rate constants. The posterior distributions for *k*_*cat*_ ratios are better defined compared to those for *k*_*cat*_ themselves. In Step 4, the catalytic rate for the dimer bound to a single substrate (DS) is faster than the rate when bound to two substrates (DSS) (Figure S22). For the representative inhibitor, both are considerably lower than the rate observed for the dimer-substrate-inhibitor complex (DSI) (Figure S14). In Step 5, for the majority of ligands in the dataset, *k*_*cat,DSI*_ is comparable to or larger than *k*_*cat,DS*_ (Figure 6). However, a few ligands (ASAP-00008489-001, ASAP-00011343-001, ASAP-00011513-001, ASAP-00012331-001, ASAP-00012335-001, ASAP-00013263-001, ASAP-00013299-001, ASAP-00013301-001, ASAP-00013407-001, ASAP-00013412-001, ASAP-00014551-001, ASAP-00014717-001, ASAP-00014750-001, ASAP-00014776-001, ASAP-00015517-001) clearly have low *k*_*cat,DSI*_.

**Figure 6.**
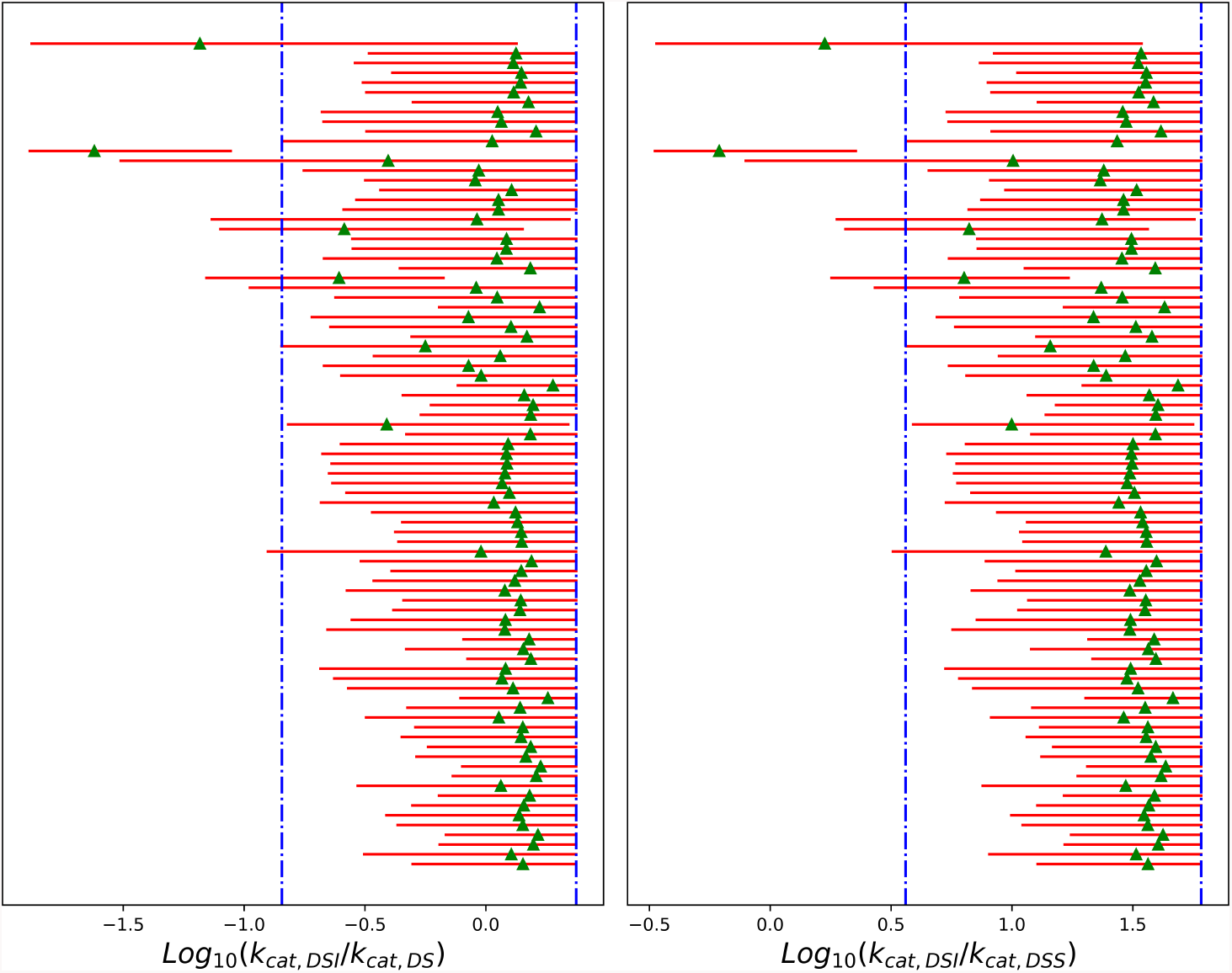
Ratios of rate constants for all ligands from the ESI4c data set. Estimates were based on 1,000 MCMC samples generated from the Bayesian posterior. Red bars represent 95% HDIs, while green triangles mark the median of the posteriors. The vertical blue lines represented for the prior distributions.

Enzyme concentration parameters are consistent with stated values except at the highest enzyme concentration (Figure 7). In Steps 1 and 2, there is still significant uncertainty in the enzyme concentrations; marginal distributions are broad. In Steps 3 and 4, the 95% HDI of enzyme concentration parameters included the stated concentrations of 25 and 50 *nM*, but the median enzyme concentration parameter for 100 *nM* is only 59.7 *nM*. It is possible that at high concentration, the enzyme precipitates or forms inactive higher-order oligomers, reducing the concentration of active enzyme.

**Figure 7.**
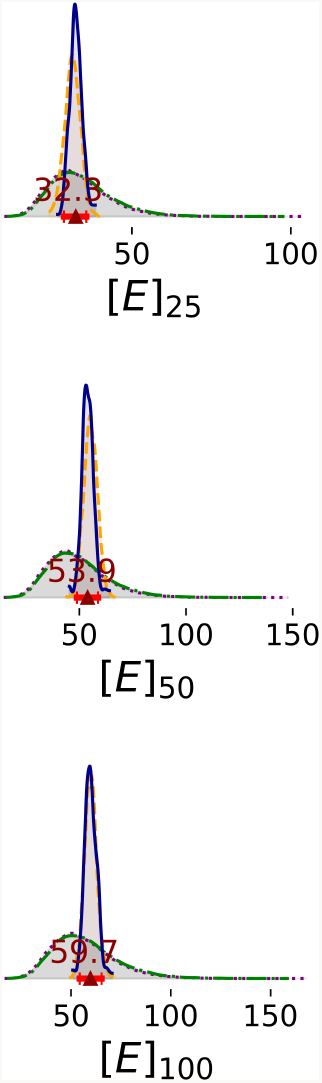
1D marginal distributions of enzyme concentrations (nM). Estimates were estimated based on 1,000 MCMC samples generated from the Bayesian posterior. Annotations are analogous to Figure 3.

### 3.2 The enzyme kinetics model closely fits nearly all data

In Step 4, the enzyme kinetics model is a close fit to nearly all of the data from the ES, ESI4c, and three ESI1c datasets (Figure 8). For some ligands, the velocity is higher than the model near the peak velocity and at the highest ligand concentration of 99.5 *µ*M. For high concentrations, ligands may have solubility limits that reduce the amount of ligand in solution. Similar results are achieved in Step 5 (Figure S23). A small subset of compounds (ASAP-0000219-001, ASAP-0000375-001, ASAP-0010712-001, ASAP-0013397-001, ASAP-0013405-001, ASAP-0013407-001, ASAP-0013423-001) exhibit a significantly broader 95% posterior predictive interval of the velocity.

**Figure 8.**
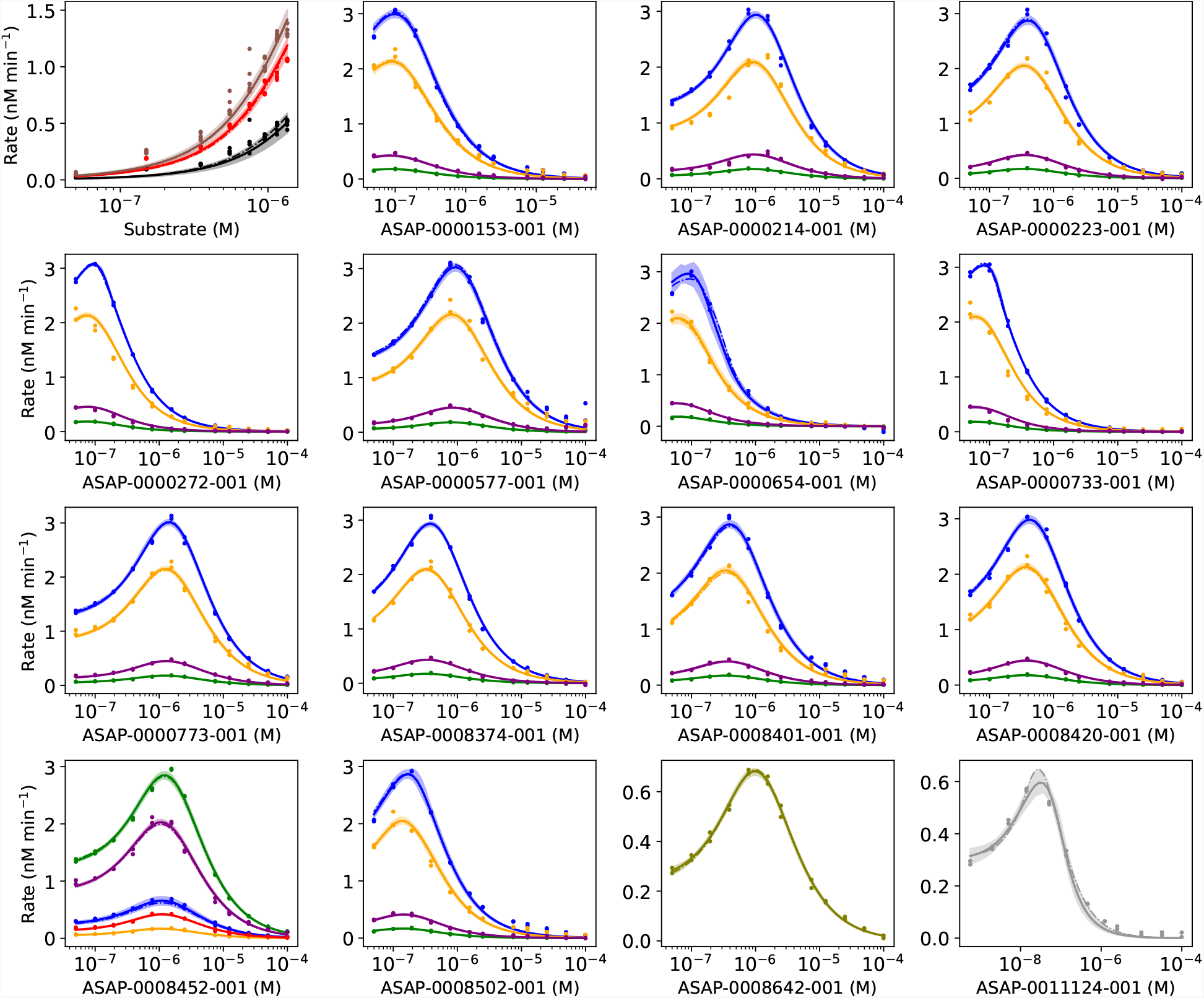
Fit of the model to ES + all ESI4c + 3 ESI1c datasets. X axes are concentrations (M). Dots are observed velocities. Velocities predicted by the model are 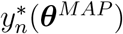 (dashed line), where ***θ***^*MAP*^ is the MAP estimate, the mean of the posterior prediction (solid line), and the 95% posterior predictive interval (shaded region). Data are colored by plate. For the upper left plot: 100 nM (brown), 50 nM (red), and 25 nM (black) of the enzyme. For other plots: enzyme 100 nM, substrate 1350 nM (blue); enzyme 100 nM, substrate 750 nM (orange); enzyme 50 nM, substrate 150 nM (purple); enzyme 100 nM, substrate 50 nM (green); enzyme 50 nM, substrate 550 nM (pink, olive, gray).

### 3.3 MERS-CoV MPro undergoes substrate-induced dimerization

We modeled substrate-induced dimerization by calculating the effect on substrate on the amount of enzyme in the monomeric versus the dimeric form. We computed the concentrations of monomeric ([M] and [MS]) and dimeric ([D], [DS], [DSS]) species given an initial enzyme concentration of 0.3 *µ*M and substrate concentration of 600 *µ*M, similar to a previous study,^45^ using thermodynamic and kinetic parameters drawn from the posterior. Monomeric and dimeric concentrations were used to compute an *apparent* binding affinity,

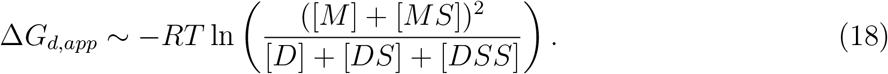

The 95% HDI of Δ*G*_*d,app*_ is [−12, −9.2] kcal/mol with a mean of −10 kcal/mol. In contrast, the 95% HDI of Δ*G*_*d*_ is [−6.7, −5.4] kcal/mol with a mean of −5.9 kcal/mol. Thus, the model predicts strong substrate-induced dimerization under conditions similar to prior work.^45^

As an orthogonal approach, we also performed mass photometry to directly monitor the oligomeric state of the enzyme (Appendix C of the Supplementary Information). Under the biochemical assay conditions, 20 nM MERS MPro is primarily monomeric and some inhibitor-induced dimerization was observed.

### 3.4 The MERS-CoV MPro dimer binds most ligands with positive cooperativity

Most inhibitors exhibited positive cooperativity with the enzyme, as the free energy difference between the second and first binding events was less than zero (Figure 9), except for ASAP-00013249-001, ASAP-00013301-001, ASAP-00013412-001, ASAP-00013894-001, and ASAP-00014900-001.

**Figure 9.**
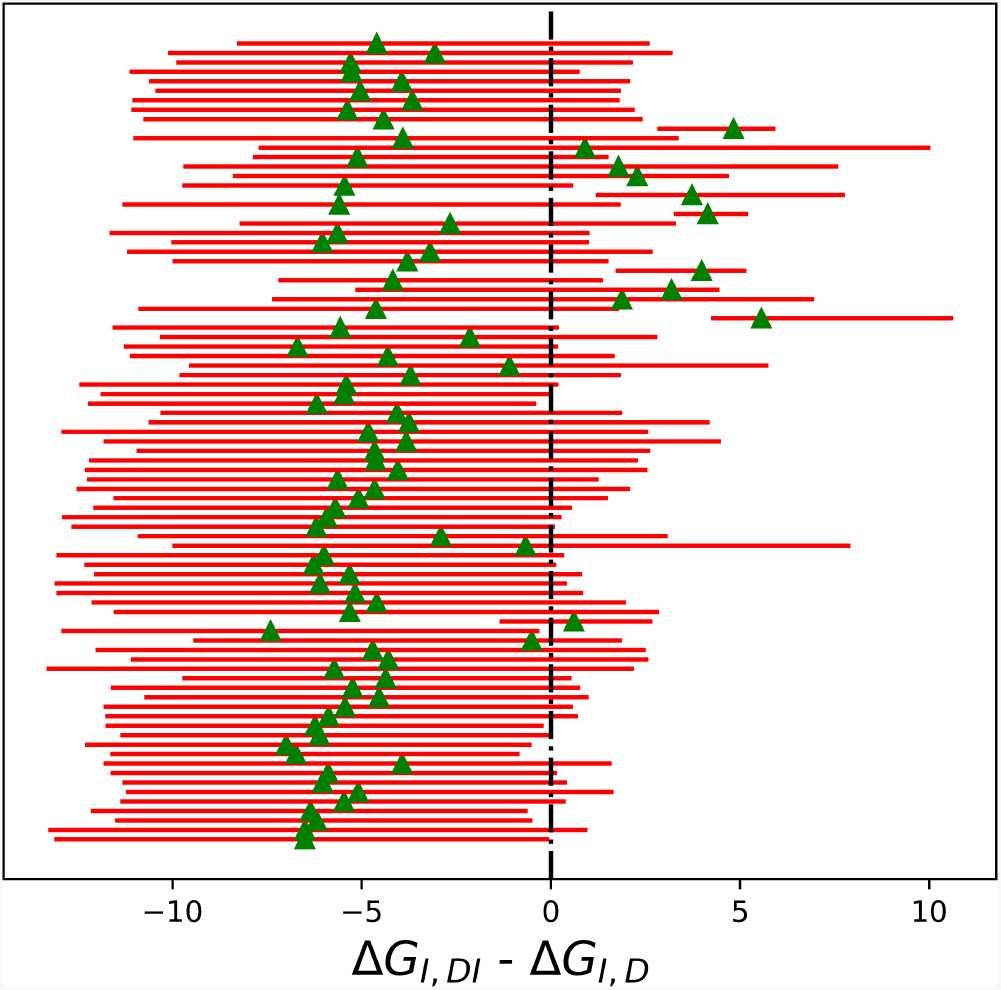
Differences in binding free energies of inhibitors (kcal/mol). Estimates were based on 1,000 MCMC samples generated from the Bayesian posterior from Step 5. Red bars represent 95% HDIs, while green triangles mark the median of the posteriors.

### 3.5 Biochemical and cellular potencies of ASAP MERS-CoV MPro inhibitors are highly correlated

Correlations between inhibition, control, and dimer pIC50s are greater than 0.8 but there are weaker correlations with cellular pEC50s (Figure 10 and Table 1). The highest correlation is observed between inhibition and control pIC50s, with the Pearson R and Spearman *ρ* above 0.93. Furthermore, the three pIC50s derived from the same biochemical CRC exhibit a high level of correlation with each other. On the other hand, the coefficients between the cellular pEC50 and the other three pIC50s are less than 0.8, indicating a clear discrepancy between the biochemical and cellular assays. This discrepancy suggests that some factors in the cellular environment, such as cellular permeability and metabolic stability, may not be captured in the biochemical assay.

**Table 1.** Correlation matrix of biochemical pIC50 and cellular pEC50 by Pearson R, Spearman *ρ*, and Kendall *τ*.

| Pearson R | Inhibition pIC50 | Control pIC50 | Dimer pIC50 |
| --- | --- | --- | --- |
| Control *pIC50* | 0.955 ± 1.493E-2 | | |
| Dimer *pIC50* | 0.888 ± 7.541E-2 | 0.878 ± 6.918E-2 | |
| Cellular *pEC50* | 0.720 ± 6.576E-2 | 0.712 ± 6.678E-2 | 0.711 ± 6.152E-2 |
| Spearman $\rho$ | Inhibition pIC50 | Control pIC50 | Dimer pIC50 |
| Control *pIC50* | 0.937 ± 2.156E-2 | | |
| Dimer *pIC50* | 0.925 ± 4.765E-2 | 0.915 ± 4.493E-2 | |
| Cellular *pEC50* | 0.677 ± 6.061E-2 | 0.659 ± 6.792E-2 | 0.718 ± 5.166E-2 |
| Kendall $\tau$ | Inhibition pIC50 | Control pIC50 | Dimer pIC50 |
| Control *pIC50* | 0.807 ± 3.352E-2 | | |
| Dimer *pIC50* | 0.839 ± 4.543E-2 | 0.805 ± 4.485E-2 | |
| Cellular *pEC50* | 0.501 ± 5.298E-2 | 0.492 ± 5.797E-2 | 0.530 ± 4.819E-2 |

**Figure 10.**
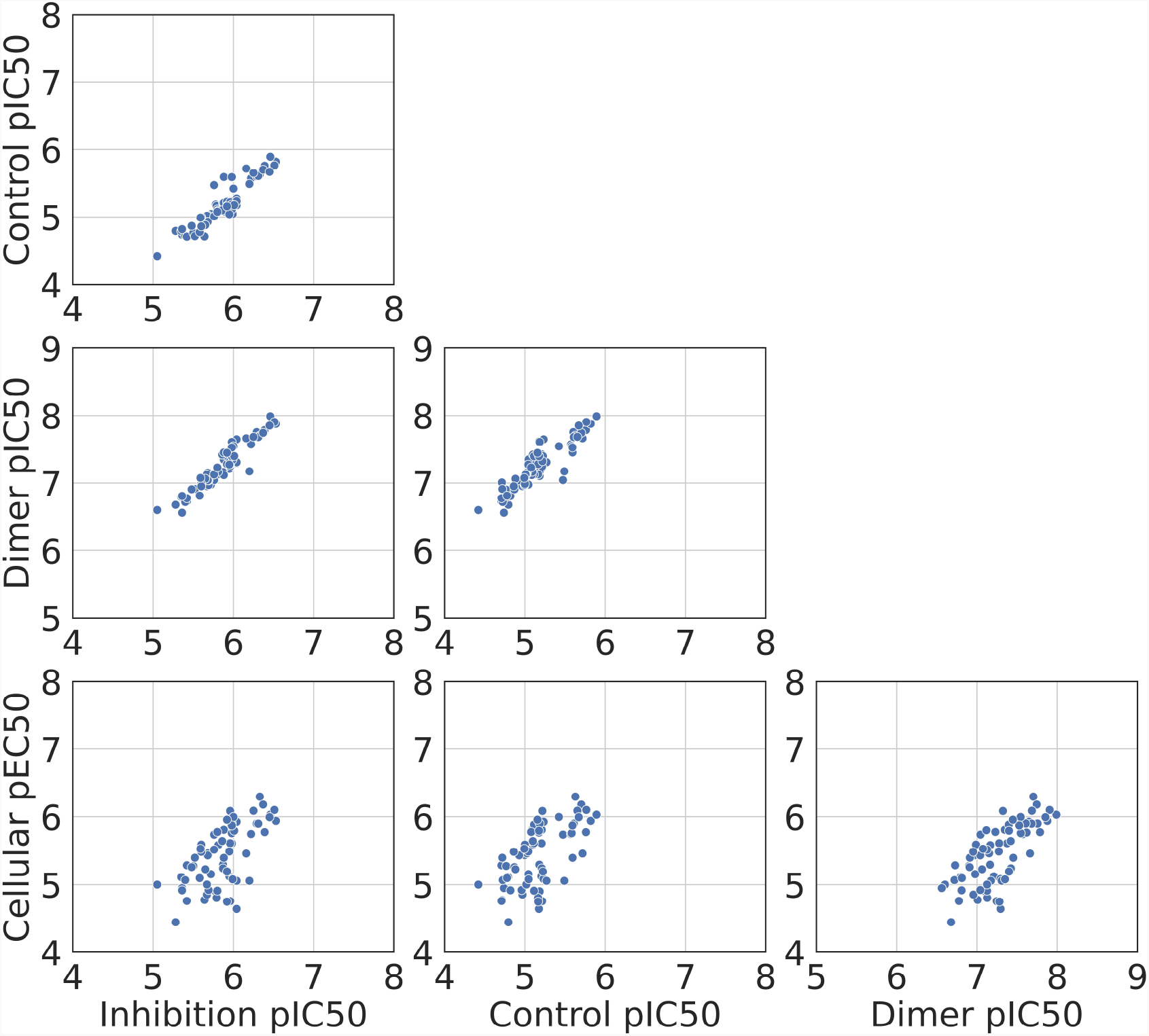
Correlogram of inhibition, control, dimer pIC50 and cellular pEC50.

The dimer pIC50 outperforms the inhibition and control pIC50 in forecasting the rank of the cellular pEC50, but has similar Pearson R and higher error (Table 1). Assuming that the statistical metrics are independent and follow a normal distribution, we evaluated p-values from the two-sample comparison of coefficients with the null hypothesis that there is no difference in the means of coefficients, then applied the Bonferroni adjustment for the comparison of 4 procedures.^46^ With the threshold at 0.05, the adjusted p-values suggested stronger evidence to reject the null hypothesis for Spearman *ρ* and Kendall *τ* (Table S3). In other words, the correlation coefficients between dimer pIC50 and cellular pEC50 are higher than the coefficients between other biochemical pIC50s and cellular pEC50 when evaluated by Spearman *ρ* and Kendall *τ*. This indicates that, according to rank order, the dimer model may align more closely with cellular responses than the inhibition and control procedures.

### 3.6 The dimer pIC90 better ranks cellular pEC90 than the dimer pIC50 ranks cellular pEC50

As many of the CRCs from antiviral assays have a Hill slope distinct from one, we hypothesized that a high level of MPro inhibition is required to improve cell viability and that biochemical pIC90 would better predict cellular pEC90 than biochemical pIC50 predicts cellular pEC50. Overall, correlation and error metrics for pIC90/pEC90 (Figure S24, Table S1, Table S2, and S4) were higher than those for pIC50/pEC50 (Figure 10, Table 1, and Table 2) when evaluated by Spearman *ρ* and Kendall *τ*.

**Table 2.**
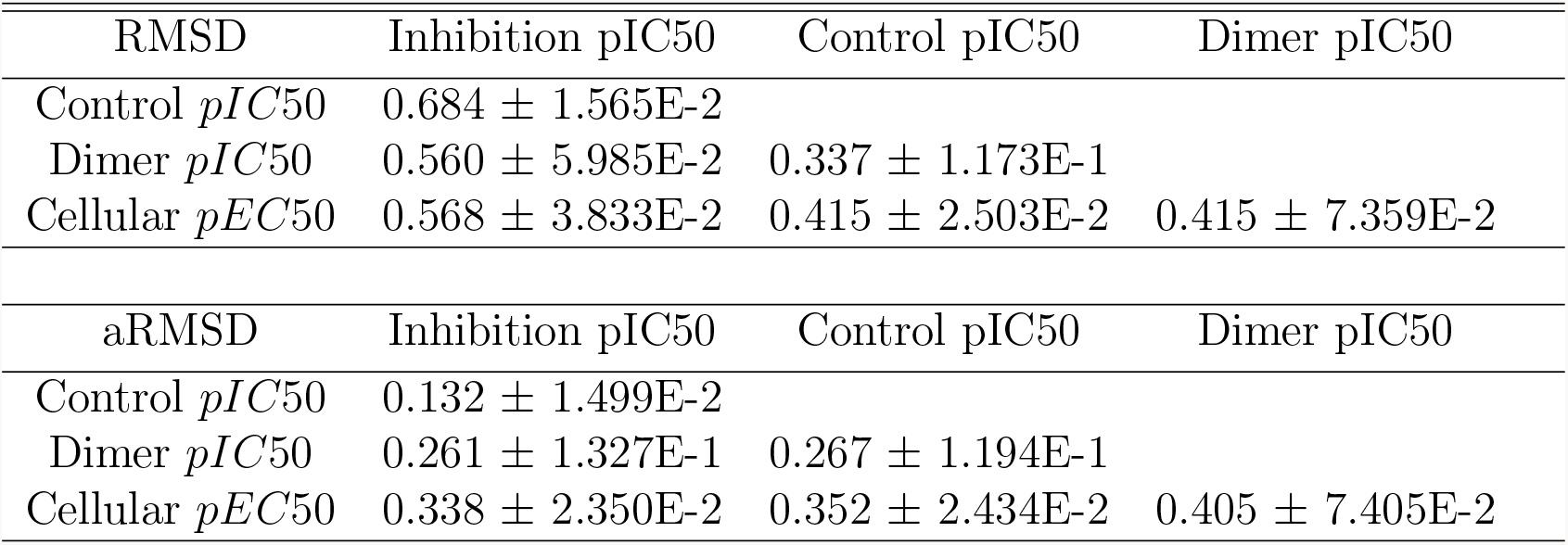
Correlation matrix of biochemical pIC50 and cellular pEC50 by RMSD and aRMSD.

## 4 Discussion

### 4.1 Enzyme kinetics experiments are sufficient to determine parameters of a complex model

We have demonstrated that enzyme kinetics experiments with varying concentrations of enzyme, substrate, and multiple inhibitors are sufficient to precisely determine most parameters of a complex enzyme kinetics model. When we recently introduced our enzyme kinetics model that incorporates both dimerization and ligand binding,^22^ we not only fit it to biochemical enzyme kinetics but also to analytical ultracentrifugation data for two variants of SARS-CoV-2 MPro: the wild-type enzyme and a mutant engineered to have a weaker dimerization affinity.^21^ Here we pursued an alternative strategy of fitting many CRCs for one enzyme.

We have also addressed the computational challenge associated with globally fitting numerous CRCs. We leveraged the structure of Bayesian posteriors to break down the fitting process into multiple steps and incrementally incorporate additional information into the model. Even with uninformative priors (besides the dimerization affinity), we observed that as the amount of data are increased from Steps 1 to 4, the binding free energies and ratios of species-specific rates are estimated more precisely. The success of global fitting extends beyond the estimation of binding affinities for the inhibitors present in the fitted datasets; the shared parameters derived from this process can also be leveraged to fit additional datasets that were not included in the original fitting procedure.

### 4.2 Parameters are within ranges reported for MERS-CoV MPro and distinct from other MPro variants

Our estimated parameters are consistent with values reported for MERS-CoV MPro enzyme kinetics. The 95% HDI of *K*_*d*_ in our paper is consistent with reported values from different measurements: 7.8 ± 0.3 *µ*M by enzymatic assay;^16^ 52 ± 5 *µ*M by analytical ultracentrifugation;^16^ and 7.7 ± 0.3 *µ*M by analytical ultracentrifugation.^45^ Based on the assumption that only dimeric enzyme is catalytically active, Tomar et al. ^16^ fit the observed rate as a function of the enzyme concentration to a model in which the apparent rate is proportional on the dimer concentration. They reported k_*cat*_ of 0.2 ± 0.02 min^−1^, which is faster than *k*_*cat,DSS*_ but slower than *k*_*cat,DS*_. Given that the apparent *k*_*cat*_ should be linear combination of turnover numbers from both species, the reported intermediate value does not raise concerns.

MERS-CoV belongs to a group of RNA viruses that have caused several human outbreaks over the past two decades, including the severe acute respiratory syndrome (SARS) CoV and SARS-CoV-2. ^4,47^ Based on phylogenetic analysis, MERS-CoV belongs to *β*-CoV lineage C and is more closely related to *Tylonycteris* bat CoV HKU4 and *Pipistrellus* bat CoV HKU5, while SARS-CoV and SARS-CoV-2 are classified into *β*-CoV lineage B.^48–50^ Our estimated *K*_*d*_ for MERS-CoV MPro is weaker than for human SARS-CoV (0.7 ± 0.02 *µ*M) ^45^ and SARS-CoV-2 (1.32 ± 0.2 *µ*M), ^21^ as well as the enzymes from closely related bat CoVs such as HKU4-CoV (0.1 ± 0.03 *µ*M) and HKU5-CoV (0.06 ± 0.01 *µ*M).^45^ The dimerization free energy is between what we reported for the SARS-CoV-2 MPro wild type (median: −8.9 kcal/mol; 95% HDI: [−9.8, −7.0] kcal/mol) and slightly stronger than the mutant (median: −4.2 kcal/mol; 95% HDI: [−4.6, −1.7] kcal/mol).^22^

Differences in dimerization affinities between MPro from different CoVs may be largely attributed to dimerization interfaces. In the case of SARS-CoV MPro, there are intermolecular polar interactions involving four amino acid pairs (Ser1-Glu166, Arg4-Glu290, Ser123-Arg298, and Ser139-Gln299). ^45^ For SARS-CoV-2 MPro, in addition to three of the pairs (Ser1-Glu166, Arg4-Glu290, and Ser139-Gln299), hydrogen bonding between the side chains of two Ser10, along with a long-distance ionic interaction between Lys12 and Glu14, were reported to contribute to this process.^51^ In contrast, only two pairs are found in MERS-CoV MPro: Ser1-Glu169 and Ser142-Gln299.^45^ The reduced number of intermolecular interactions likely cause MERS-CoV MPro to have a weaker *K*_*d*_ compared to the enzymes from the other human CoVs. On the other hand, the differences in the *K*_*d*_ values between MERS-CoV MPro and its closely related HKU4-CoV and HKU5-CoV can be attributed to the non-conserved residues located in the N-terminal finger, the N-terminal helix, and domain III.^16^

Comparing other enzyme kinetics parameters for SARS-CoV-2 MPro from our previous analysis^22^ and MERS from our present analysis illustrates variations in how the enzyme can behave and interact with ligands. The binding free energy of substrate to the monomer Δ*G*_*S,M*_ is comparable in MERS-CoV MPro (median: −5.1; 95% HDI: [−5.7, −4.4] kcal/mol) and SARS-CoV-2 MPro (median: −4.7; 95% HDI: [−4.8, −4.5] kcal/mol), but dimerization has a much greater effect on substrate binding for MERS-CoV MPro. While the binding affinity of the substrate for the dimer Δ*G*_*S,D*_ is much lower than Δ*G*_*S,M*_ for MERS-CoV MPro (median: −9.0; 95% HDI: [−9.9, −8.4] kcal/mol), it is similar to Δ*G*_*S,M*_ for SARS-CoV-2 MPro (median: −4.7; 95% HDI: [−6.0, −4.0] kcal/mol). There is also a contrast in binding cooperativity behavior; while the binding free energy of the substrate to the dimer-substrate complex Δ*G*_*S,DS*_ for MERS-CoV MPro (median −10.0; 95% HDI: [−10.1, −9.3] kcal/mol) is lower than Δ*G*_*S,D*_, indicating positive cooperativity, Δ*G*_*S,DS*_ for SARS-CoV-2 MPro (median −0.8; 95% HDI: [−2.7, 0.] kcal/mol) is higher than Δ*G*_*S,D*_, indicating negative cooperativity. Relative rate constants also differ between MPro variants. For wild-type SARS-CoV-2 MPro, all of the rates − *k*_*cat,DS*_, *k*_*cat,DSS*_, and *k*_*cat,DSI*_ - are comparable.^22^ For the mutant, *k*_*cat,DSI*_ and *k*_*cat,DSS*_ are comparable to each other and larger than *k*_*cat,DS*_. Here, we report that *k*_*cat,DSI*_ and *k*_*cat,DS*_ are comparable to each other and larger than *k*_*cat,DSS*_ for MERS-CoV MPro. These comparisons suggest that coronaviruses could employ different strategies for the regulation of enzyme activity. Compared to SARS-CoV-2 MPro, MERS-CoV MPro requires more enzyme to dimerize and become active, but the dimer binds more tightly to the substrate.

As a word of caution, differences in enzyme kinetics parameters may not be due the proteins themselves, but can be affected by assay conditions such as the choice of substrate. Our study used the substrate [5-FAM]-AVLQSGFR-[Lys(Dabcyl)]-K-amide. In contrast, Nashed et al. ^21^ used Dabsyl-KTSAVLQ/SGFRKM-E(Edans)-NH2, a longer peptide that could have greater difficulty occupying both binding sites. Additional data and analyses would be required to dissect whether parameter differences originate from the protein sequence or the assay conditions.

### 4.3 Full CRC fitting and dimer pIC90 calculations are recommended for drug discovery targeting MERS-CoV MPro and similar enzymes

We recommend interpreting CRCs of MERS-CoV enzyme inhibition by fitting an enzyme kinetics model and calculating dimer pIC90s. We compared three data analysis procedures using Pearson R, Spearman *ρ*, and Kendall *τ* correlation metrics based on two assumptions: the null hypothesis for multiple-sample comparisons that there are no differences in the means of the coefficients; and the correlation coefficients follow a normal distribution. While bounded metrics cannot strictly follow a normal distribution, these assumptions are a reasonable approximation. Based on these assumptions, all three data analysis procedures that we evaluated yielded inhibition constants that are correlated with cellular efficacy. While all three biochemical procedures have similar Pearson R, dimer pIC50/90 generally have a higher Spearman *ρ* and Kendall *τ* rank correlation with cellular pEC50/90. Additionally, the pIC90 was found to be more predictive than the pIC50. In the context of a drug discovery campaign, the major objective of performing biochemical assays is to prioritize a subset of compounds to test in more expensive and challenging antiviral assays. Rank order correlations are the most relevant metrics for prioritization decisions.

Beyond achieving the highest rank correlations with cellular efficacy, fitting the enzyme kinetics model provides insight into the mechanism of specific inhibitors. MPro enzyme kinetics can be altered by inhibitor binding free energies to different species including the monomer, dimer, and dimer-ligand complexes. Determining these parameters can help evaluate whether a compound promotes dimerization or binds cooperativity to the target. The parameters can also quantify whether inhibitor binding to one catalytic site increases activity of the opposite site. Finally, determined parameters can be used to validate results from molecular simulations. Because molecular simulations are performed with specific species, inhibitor binding free energies to specific species can be directly compared to forecasts from these calculations. Parameters from our fitting procedure have been used as benchmarks to validate an ASAP computational workflow Castellanos et al. ^52^ and in a blind challenge for the computational chemistry community MacDermott-Opeskin et al. ^53^.

The main drawback of full CRC fitting is its relative complexity. Fortunately, we have made our code freely available.^44^ In service of the ASAP drug discovery campaign targeting SARS-CoV-2/MERS-CoV MPro, the procedure was integrated into an automated data analysis pipeline that presents results on CDD Vault.^38^

Beyond the present application to steady-state kinetics of MPro inhibited by noncovalent and reversible covalent inhibitors, our steady-state enzyme kinetics model^22^ is a natural starting point for time-dependent extensions and applications to other enzymes. Our model can be extended to irreversible covalent inhibitors via a time-dependent scheme that separates reversible recognition from covalent inactivation (e.g. reporting kinetic efficiency through *k*_inact_*/K*_*I*_ and fitting progress curves or time-staggered CRCs). More broadly, hysteresis - potentially arising from slow conformational adaptation or from oligomerization equilibria such as ligand- or substrate-induced dimerization^54^ - can be captured by augmenting the model with explicit exchange rates. Besides MERS-CoV MPro, substrate/ligand-induced dimerization has been observed in multiple serine proteases.^55,56^ Biphasic CRCs have been reported for the cysteine protease caspase-1,^57^ and may be a factor in other enzymes.

## 5 Conclusion

We have developed a multi-step statistical analysis procedure to fit an enzyme kinetics model that incorporates dimerization and ligand binding to many CRCs with multiple inhibitors from a drug discovery campaign. The analysis precisely determines many binding parameters and ratios of rate constants and quantifies substrate-induced dimerization and ligand binding cooperativity. For the leads in the ASAP drug discovery campaign targeting MERS-CoV and SARS-CoV-2, inhibition constants from multiple data analysis procedures are highly correlated with cellular efficacy. pIC90s estimated by simulating CRCs at high enzyme concentration are have higher rank correlation with cellular pEC90 than the other tested procedures. Code implementing the procedure is freely available and is recommended for the interpretation of biphasic CRCs from MERS-CoV MPro and enzymes with similar properties.

## Supporting information

Normalized concentration response curves

## Acknowledgement

This project was supported in part by NIAID of the National Institutes of Health under award no. U19AI171399 (JDC) and the NIH/NCI Cancer Center Support Grant P30 CA008748 (JDC).

This work used the Jetstream2 cloud-based environment at Indiana University through allocation MCB150144 from the Advanced Cyberinfrastructure Coordination Ecosystem: Services & Support (ACCESS) program, which is supported by National Science Foundation grants #2138259, #2138286, #2138307, #2137603, and #2138296.

The content is solely the responsibility of the authors and does not necessarily represent the official views of the National Science Foundation or the National Institutes of Health.

## Supporting Information Available

Extended methodological analyses and validation, including: steady-state assessment of enzyme assays; convergence diagnostics for Bayesian MCMC sampling; posterior marginal and joint distributions for binding free energies, rate constants, and enzyme concentrations across ES, ESI4c, and ESI1c datasets; model fits to biochemical inhibition datasets; correlation analyses linking biochemical and cellular potency metrics; and mass photometry analysis of MERS-CoV MPro oligomerization.

