## Appendices and Supplementary Figures and Tables for "Linking biochemical and cellular efficacy of MERS coronavirus main protease inhibitors"

### Supplementary Information for “Linking biochemical and cellular efficacy of MERS coronavirus main protease inhibitors”

### Contents

**Appendix A. Enzyme catalysis was in a steady state.** **5**

Figure S1. Example fluorescence time courses with short- and long-window linear fits. . . . . . . . . . . . . . . . . . . . . . . . . . . . . . . . . . . . . . . 5

Figure S2. Plate-wide comparison of 10-min fit extrapolation to 1-hr measurements. 5

**Appendix B. Bayesian credible intervals were converged.** **6**

Figure S3. Convergence of percentiles of the Bayesian posterior for fitting ES datasets 6

**Figure S7. 1D marginal distributions for ES fitting of MERS-CoV MPro** **14**

**Figure S8. 2D joint marginal distributions for pairs of binding free energies of fititng ES dataset of MERS-CoV MPro** **15**

**Figure S9. Fit of the model to ES and ESI4c datasets for ASAP-0000214** **16**

**Figure S10. 1D marginal distributions for binding free energies of fitting ES and ASAP-0000214 dataset of MERS-CoV MPro** **17**

**Figure S11. 2D joint marginal distributions for pairs of binding free energies of fititng ES and ASAP-0000214 dataset of MERS-CoV MPro** **18**

**Figure S12. Heat map of the correlation matrix estimated from the Bayesian posterior for free energies of fitting ES and ASAP-0000214 datasets of**

**MERS-CoV MPro** **19**

**Figure S13. Linear correlation between $\Delta GK_{I,D}$, $\Delta G_{S,DI}$, and $\Delta G_{I,DI}$ of fitting ES and ASAP-0000214 datasets MERS-CoV MPro.** **19**

**Figure S14. 1D marginal distributions for rate constants of fitting ES and ASAP-0000214 datasets of MERS-CoV MPro** **20**

**Figure S15. 2D marginal distributions for pairs of rate constants of fitting ES and ASAP-0000214 datasets of MERS-CoV MPro** **20**

**Figure S16. 1D marginal distributions for enzyme concentrations of fitting ES and ASAP-0000214 datasets of MERS-CoV MPro** **20**

**Figure S17. 2D joint marginal distributions for pairs of some binding free energies of fitting ES and all ESI4c datasets** **22**

**Figure S18. Heat map of the correlation matrix estimated from the Bayesian posterior for some binding free energies of fitting ES and all ESI4c datasets** **22**

**Figure S19. Fit of the model to ES and all ESI4c datasets** **22**

**Figure S20. 2D joint marginal distributions for pairs of some binding free energies of fitting ES, all ESI4c, and 3 ESI1c datasets** **24**

**Figure S21. Heat map of the correlation matrix estimated from the Bayesian posterior for some binding free energies for fitting ES, all ESI4c, and 3 ESI1c datasets** **24**

**Figure S22. Ratios of rate constants of MERS-CoV MPro** **24**

**Figure S23. Fit of the model to ESI1c datasets** **26**

**Figure S24. Correlogram of inhibition, control, dimer pIC90 and cellular pEC90** **32**

**Appendix C. MPro is primarily monomeric.** **33**

Figure S25. Oligomerization of MERS MPro monitored by mass photometry. . . . 33

**Table S1. Correlation matrix of biochemical pIC90 and cellular pEC90 by Pearson R, Spearman $\rho$, and Kendall $\tau$** **34**

**Table S2. Correlation matrix of biochemical pIC90 and cellular pEC90 by RMSD and aRMSD** **34**

**Table S3. p-value in comparison of correlation for $pIC50$s** **35**

**Table S4. p-value in comparison of correlation for $pIC90$s** **36**

### Appendix A. Enzyme catalysis was in a steady state.

Fluorescence time courses are largely linear over the measurement window, consistent with approximately steady-state behavior (Figure S1). Across most examples, the apparent rate slows slightly over time (the 0–10 min fit is marginally steeper than the 0–60 min fit), while in a few cases the trajectory slightly accelerates (the long-window fit is steeper or the points bend upward). These modest departures from perfect linearity are consistent with common time-dependent effects in enzyme assays, including substrate depletion (leading to slowing) or product:enzyme interactions (e.g., product inhibition or activation), altering the effective rate over time.

#### Figure S1. Example fluorescence time courses with short- and long-window linear fits.

Raw fluorescence (RFU) versus time is shown for six representative wells (A3, L3, M3, H4, I4, and D1) from a ESI4c plate with an enzyme concentration of 100 nM and substrate concentration of 1350 nM. Points indicate the measured fluorescence at each acquisition time. Solid lines show the least-squares linear fit over the first 10 minutes (0–600 s), and dashed lines show the least-squares linear fit over a 1 hour window (0–3600 s). Panel titles report well, inhibitor identity, and inhibitor concentration. Vertical guide lines indicate 10 min and 60 min.

In all four plates of the ESI4c dataset, the 1-hour endpoint fluorescence is largely consistent with the initial velocity estimated from the first 10 minutes (Figure S2). Across all wells within each plate, extrapolating the 0–10 min fit to 1 hour produces predicted RFU values that fall close to the identity line when compared to the measured 1-hour fluorescence, with high Pearson and Spearman correlations reported in each panel. Together, these results support using the 1-hour measurement as a reliable summary readout (as we have done for ESI1c) that remains strongly aligned with the short-window, initial-rate regime.

**Figure S2. Plate-wide comparison of 10-min fit extrapolation to 1-hr measurements.**

### Appendix B. Bayesian credible intervals were converged

The convergence of sampling from Bayesian posteriors was evaluated based on the 5-th, 25-th, 50-th, 75-th and 95-th percentiles of the marginal probability of parameters. In representatives of all analyses (Figure S3, S4, S5, S6), these percentiles exhibited minimal changes as the number of samples increases. Estimated standard errors were negligible. This convergence indicates that the posterior distributions have been thoroughly sampled after a small number of samples from the posterior.

**Figure S3. Convergence of percentiles of the Bayesian posterior for fitting ES datasets**

10,000 samples were drawn from the Bayesian posterior using the NUTS sampler. Lines correspond to the 5-th (blue circle), 25-th (green square), 50-th (red diamond), 75-th (cyan upward triangle) and 95-th (magenta downward triangle) percentile. The error bars, which are too small to be visible, are standard deviations estimated by 100 bootstrapping samples.

Example well traces

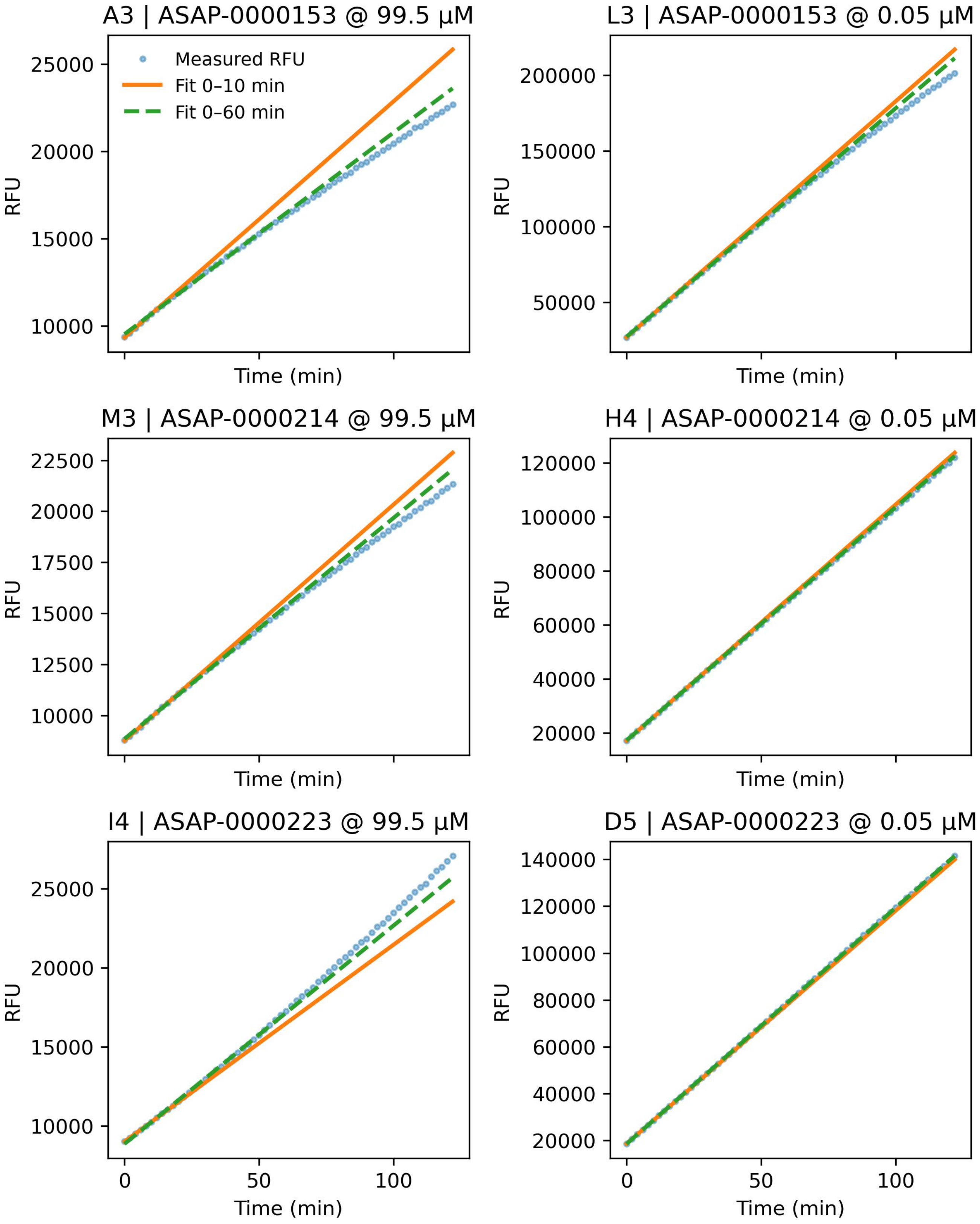

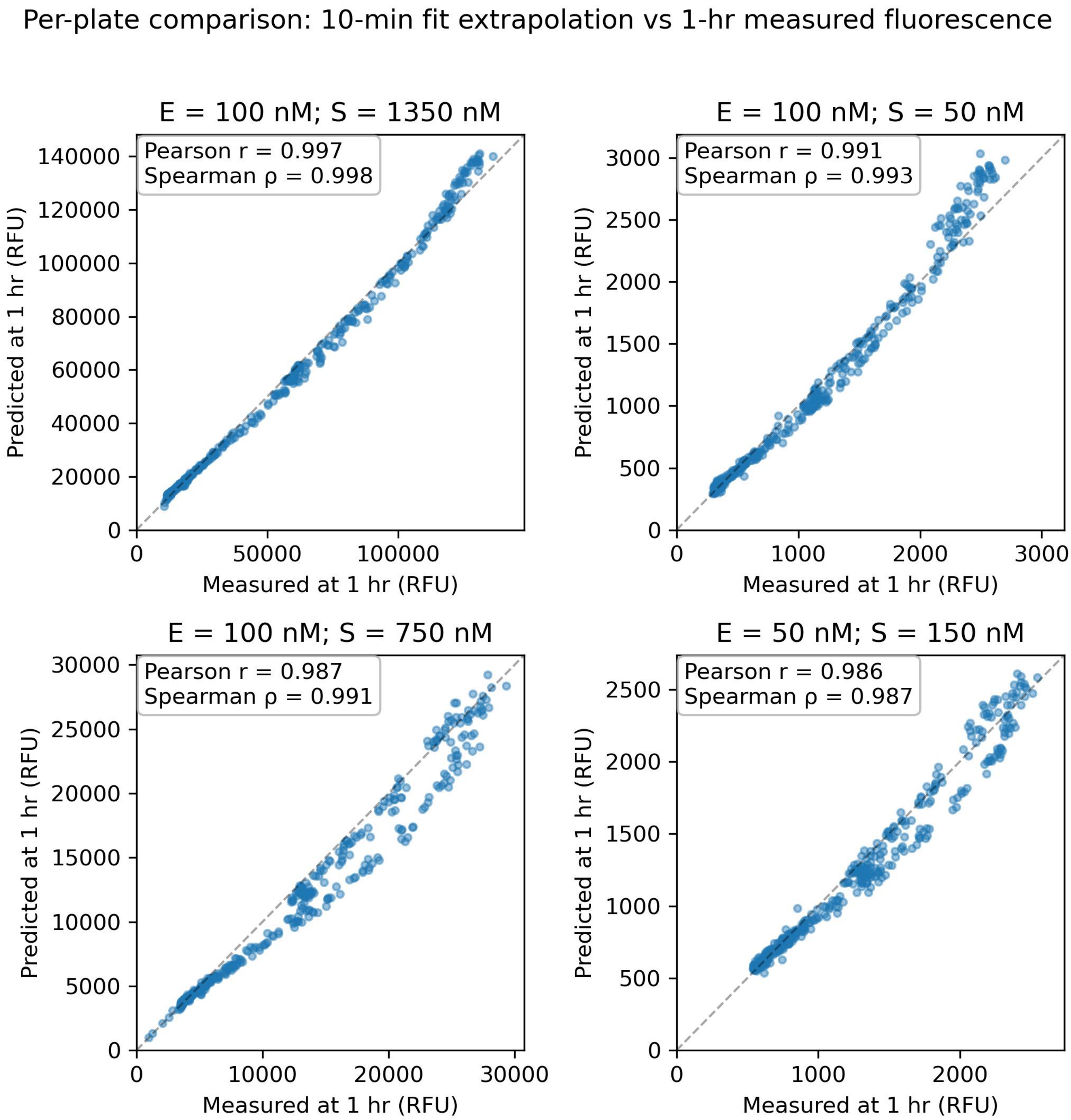
Per-plate comparison: 10-min fit extrapolation vs 1-hr measured fluorescence
E = 100 nM; S = 1350 nM
Pearson r = 0.997
Spearman ρ = 0.998
E = 100 nM; S = 50 nM
Pearson r = 0.991
Spearman ρ = 0.993
E = 100 nM; S = 750 nM
Pearson r = 0.987
Spearman ρ = 0.991
E = 50 nM; S = 150 nM
Pearson r = 0.986
Spearman ρ = 0.987
Predicted at 1 hr (RFU)
Measured at 1 hr (RFU)

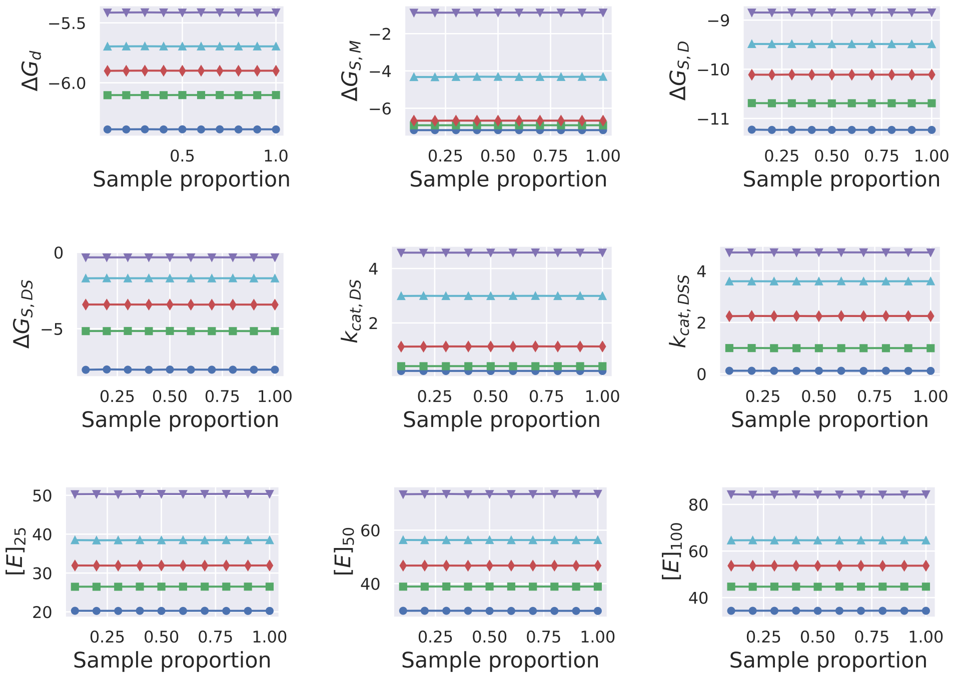

$\Delta G_d$
Sample proportion
$\Delta G_{S,M}$
Sample proportion
$\Delta G_{S,D}$
Sample proportion
$\Delta G_{S,DS}$
Sample proportion
$k_{cat,DS}$
Sample proportion
$k_{cat,DSS}$
Sample proportion
$[E]_{25}$
Sample proportion
$[E]_{50}$
Sample proportion
$[E]_{100}$
Sample proportion

#### Figure S4. Convergence of percentiles of the Bayesian posterior of key parameters for fitting ES and ASAP-0000214 datasets

10,000 samples were drawn from the Bayesian posterior using the NUTS sampler. Lines correspond to the 5-th (blue circle), 25-th (green square), 50-th (red diamond), 75-th (cyan upward triangle) and 95-th (magenta downward triangle) percentile. The error bars, which are too small to be visible, are standard deviations estimated by 100 bootstrapping samples.

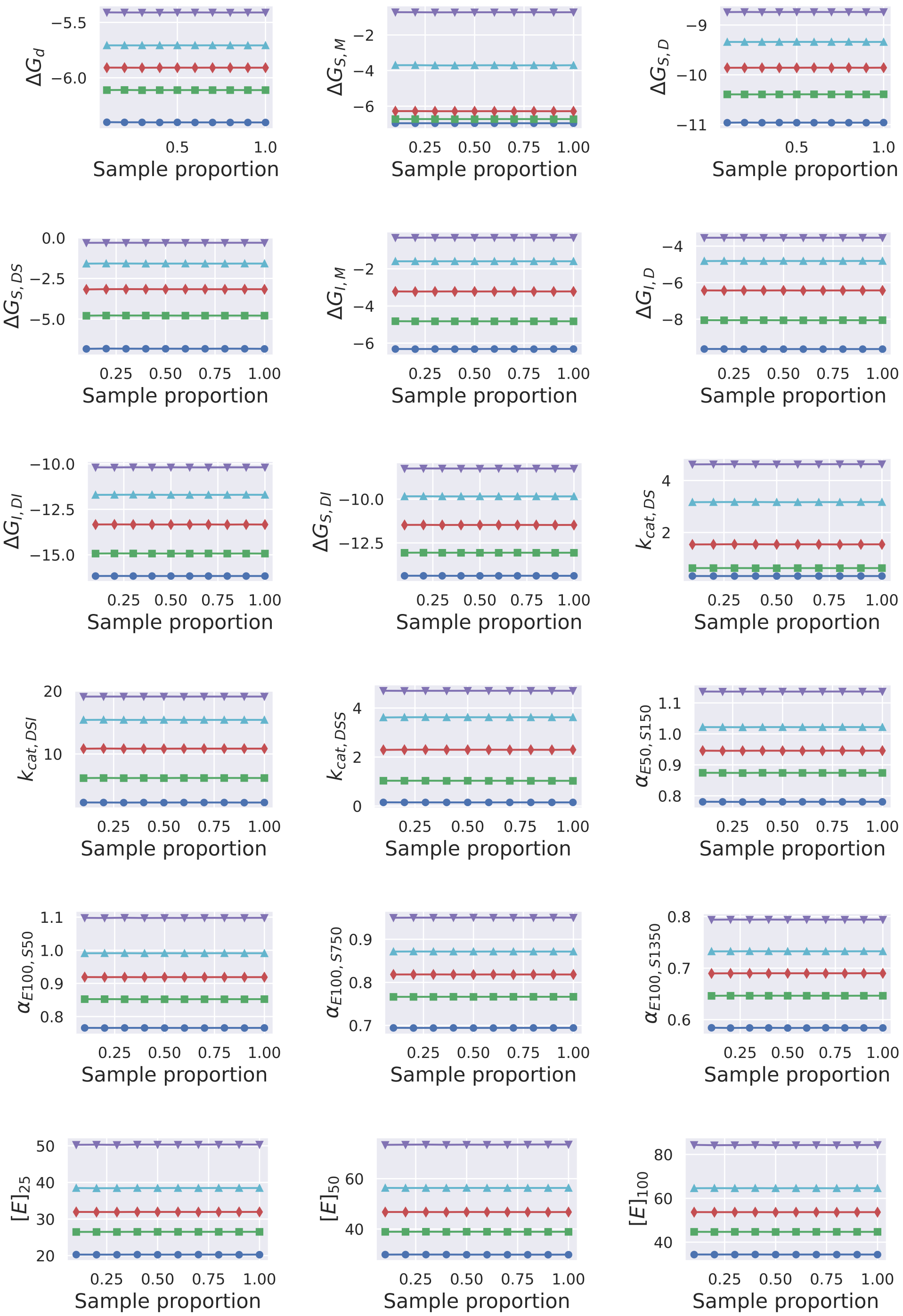

$\Delta G_d$
$\Delta G_{S,M}$
$\Delta G_{S,D}$
$\Delta G_{S,DS}$
$\Delta G_{I,M}$
$\Delta G_{I,D}$
$\Delta G_{I,DI}$
$\Delta G_{S,DI}$
$k_{cat,DS}$
$k_{cat,DSI}$
$k_{cat,DSS}$
$\alpha_{E50,S150}$
$\alpha_{E100,S50}$
$\alpha_{E100,S750}$
$\alpha_{E100,S1350}$
$[E]_{25}$
$[E]_{50}$
$[E]_{100}$
Sample proportion

#### Figure S5. Convergence of percentiles of the Bayesian posterior for shared parameters for fitting ES, all ESI4c, and 3 ESI1c datasets

1,000 samples were drawn from the Bayesian posterior using the NUTS sampler. Lines correspond to the 5-th (blue circle), 25-th (green square), 50-th (red diamond), 75-th (cyan upward triangle) and 95-th (magenta downward triangle) percentile. The error bars, which are too small to be visible, are standard deviations estimated by 100 bootstrapping samples.

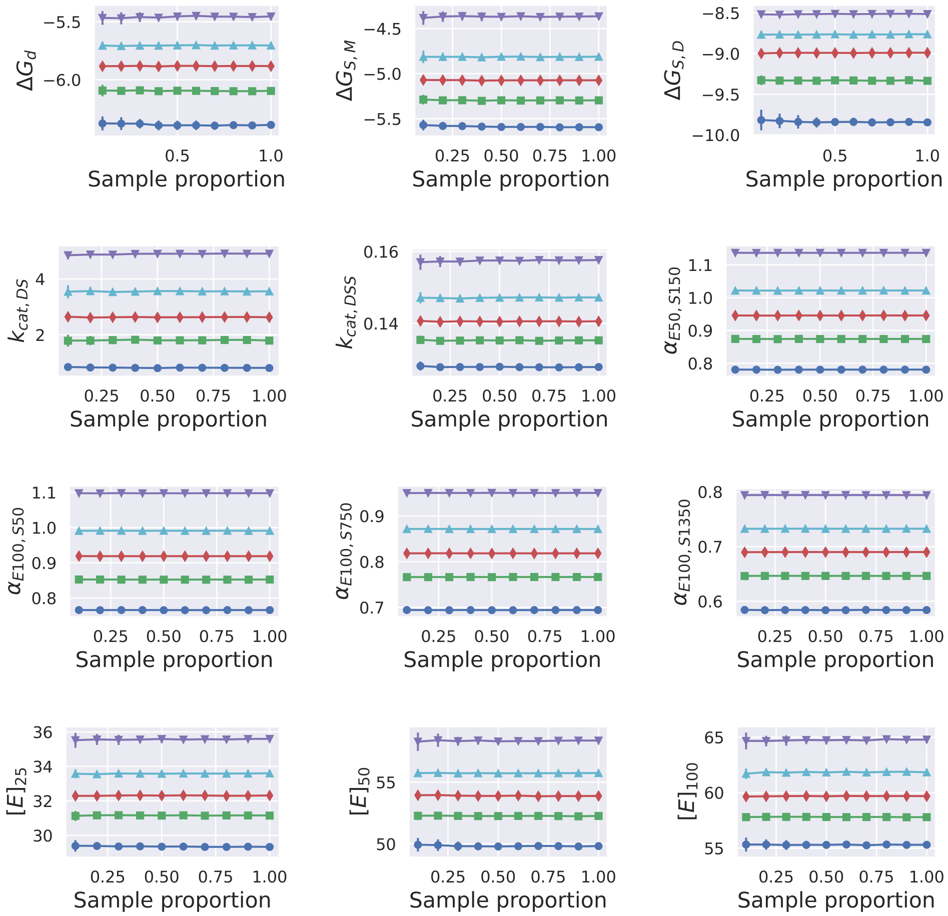

#### Figure S6. Convergence of percentiles of the Bayesian posterior for key parameters for fitting one ESI1c dataset

1,000 samples were drawn from the Bayesian posterior using the NUTS sampler. Lines correspond to the 5-th (blue circle), 25-th (green square), 50-th (red diamond), 75-th (cyan upward triangle) and 95-th (magenta downward triangle) percentile. The error bars, which are too small to be visible, are standard deviations estimated by 100 bootstrapping samples.

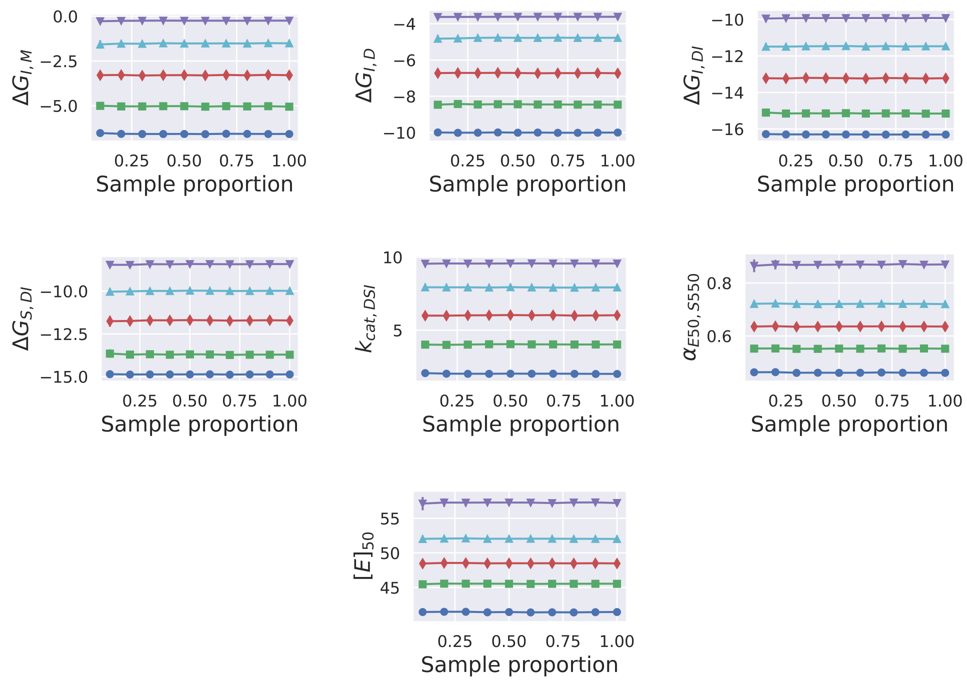

### Fitting of ES datasets

#### Figure S7. Representative 1D marginal distributions for ES fitting of MERS-CoV MPro

1D marginal probability densities for parameters were estimated based on 10,000 MCMC samples generated from the Bayesian posterior for all datasets. Red bars represent 95% HDIs. The green triangle marks the median of the posterior.

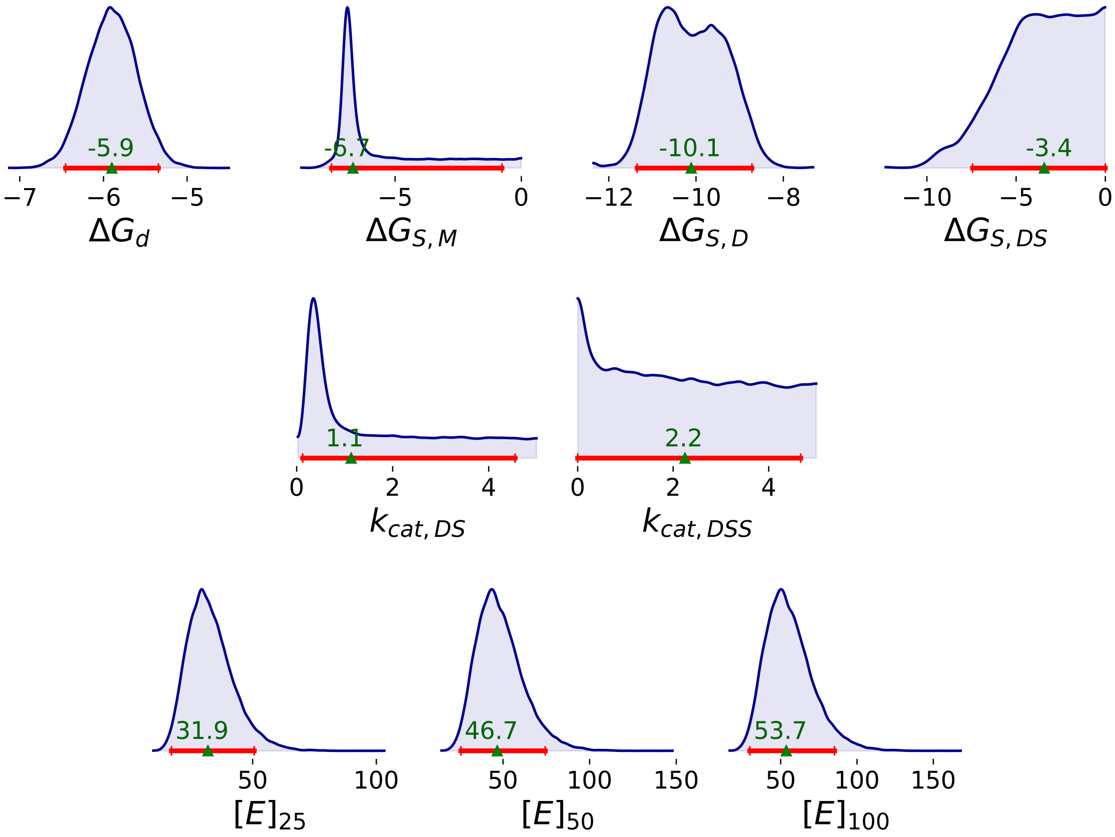

#### Figure S8. 2D joint marginal distributions for pairs of binding free energies of fitting ES datasets of MERS-CoV MPro

2D joint marginal probability densities were estimated based on 10,000 MCMC samples generated from the Bayesian posterior for all datasets.

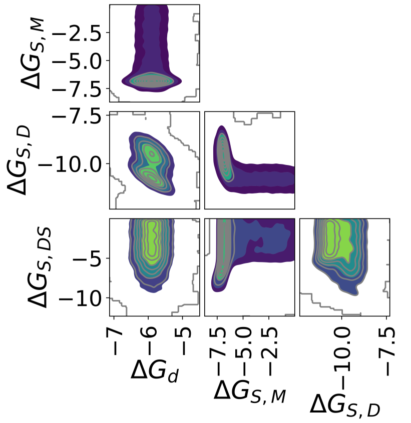

### Fitting of ES + one ESI4c datasets

#### Figure S9. Fit of the model to ES and ESI4c datasets for ASAP-0000214

Dots are the observed response. X axes are concentrations (M). The theoretical response $y_n * (\boldsymbol{\theta}^{MAP})$ is represented by the dashed line, where $\boldsymbol{\theta}^{MAP}$ is the MAP estimate, the mean of the posterior prediction is solid line, and the 95% posterior predictive interval is the shaded region. Figure a: 100 nM (brown), 50 nM (red), and 25 nM (black) of the enzyme. Figure b: enzyme 100 nM, substrate 1350 nM (blue); enzyme 100 nM, substrate 750 nM (orange); enzyme 50 nM, substrate 150 nM (purple); enzyme 100 nM, substrate 50 nM (green); enzyme 50 nM, substrate 550 nM.

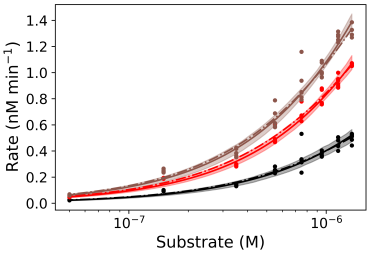

(a)

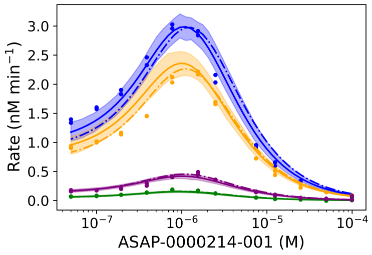

(b)

#### Figure S10. 1D marginal distributions for binding free energies of fitting ES and ASAP-0000214 datasets of MERS-CoV MPro.

1D marginal probability densities for binding free energies were estimated based on 10,000 MCMC samples generated from the Bayesian posterior for all datasets. Red lines represent 95% HDIs. The green triangle marks the median of the posterior.

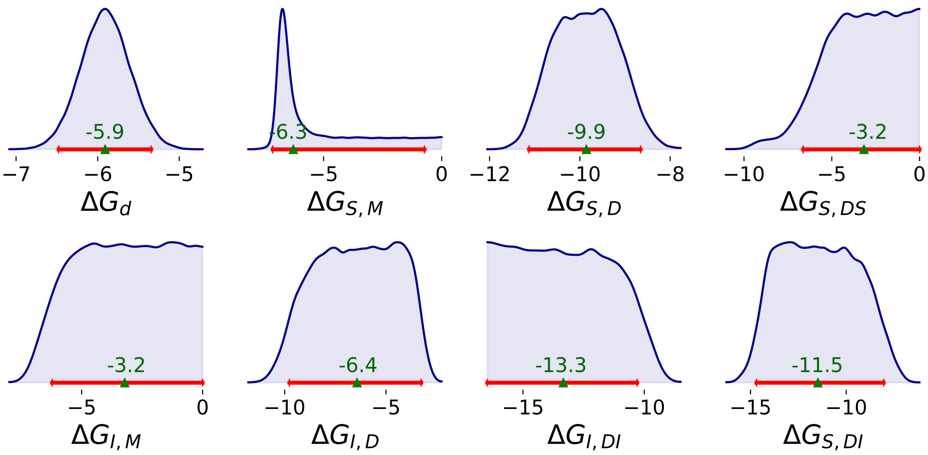

#### Figure S11. 2D joint marginal distributions for pairs of binding free energies of fitting ES and ASAP-0000214 datasets of MERS-CoV MPro

2D joint marginal probability densities were estimated based on 10,000 MCMC samples generated from the Bayesian posterior for all datasets.

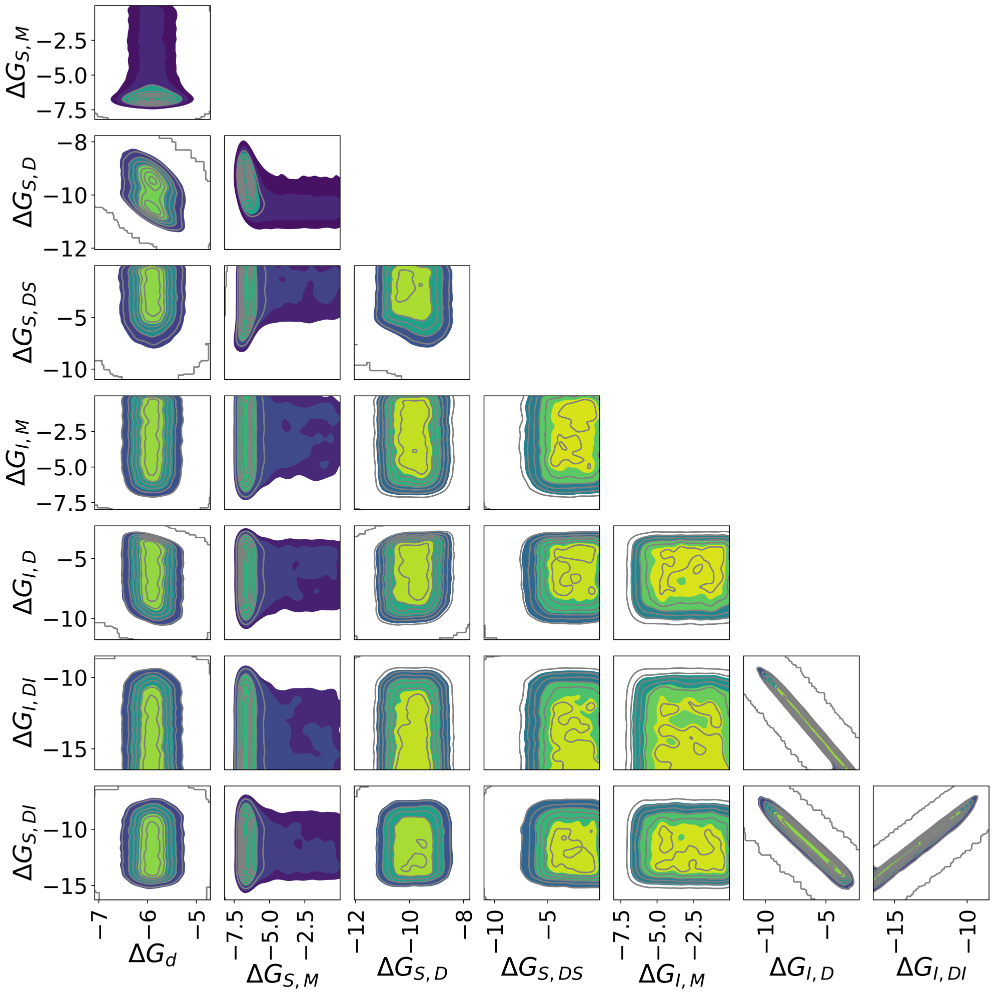

**Figure S12. Heat map of the correlation matrix estimated from the Bayesian posterior for binding free energies of fitting ES and ASAP-0000214 datasets of MERS-CoV MPro**

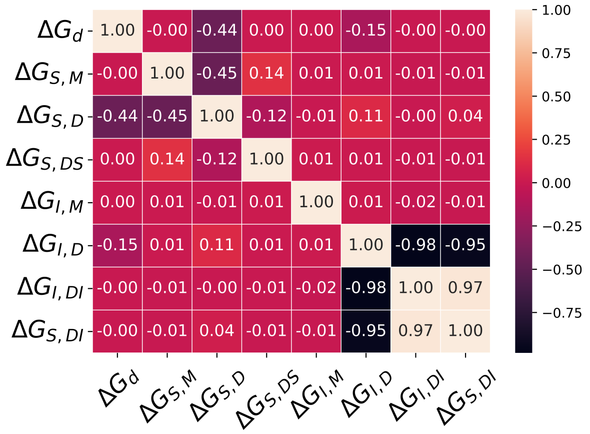

**Figure S13. Linear correlation between log$K_{I,D}$, log$K_{S,DI}$, and log$K_{I,DI}$ of fitting ES and ASAP-0000214 datasets of MERS-CoV MPro.**

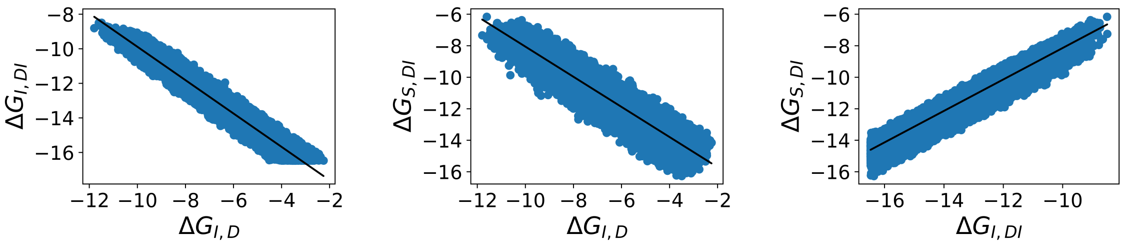

(a) (b) (c)

#### Figure S14. 1D marginal distributions for rate constants of fitting ES and ASAP-0000214 datasets of MERS-CoV MPro

1D marginal probability densities for kinetic parameters were estimated based on 10,000 MCMC samples generated from the Bayesian posterior for all datasets. Red lines represent 95% HDIs. The green triangle marks the median of the posterior.

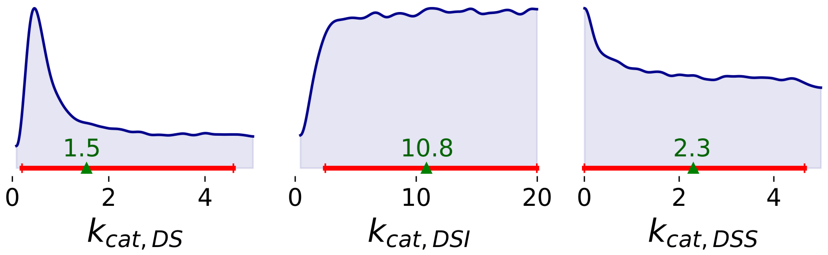

#### Figure S15. 2D marginal distributions for pairs of rate constants of fitting ES and ASAP-0000214 datasets of MERS-CoV MPro

2D joint marginal probability densities were estimated based on 10,000 MCMC samples generated from the Bayesian posteriors.

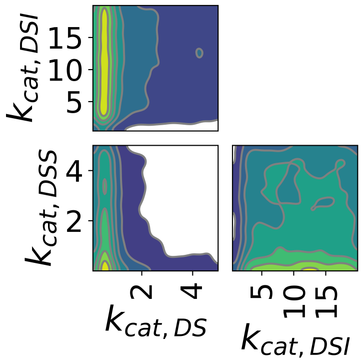

#### Figure S16. Representative 1D marginal distributions for enzyme concentrations of fitting ES and ASAP-0000214 datasets of MERS-CoV MPro

1D marginal probability densities were estimated based on 10,000 MCMC samples generated from the Bayesian posterior for all datasets. Red lines represent 95% HDIs. The green triangle marks the median of the posterior.

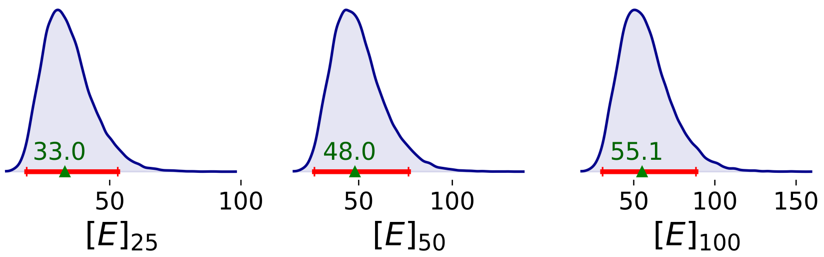

### Fitting of ES and all ESI4c datasets

#### Figure S17. 2D joint marginal distributions for pairs of some binding free energies of fitting ES and all ESI4c datasets

2D marginal probability densitie were estimated based on 1,000 MCMC samples generated from the Bayesian posterior for fitting ES and all ESI4c datasets.

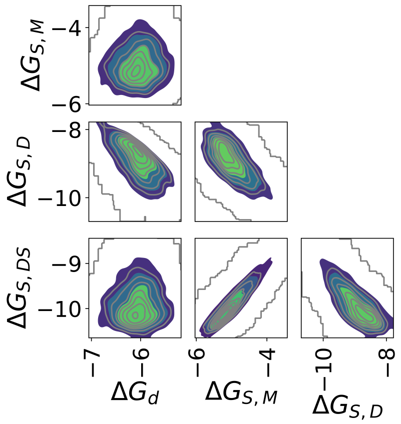

#### Figure S18. Heat map of the correlation matrix estimated from the Bayesian posterior for some binding free energies of fitting ES and all ESI4c datasets

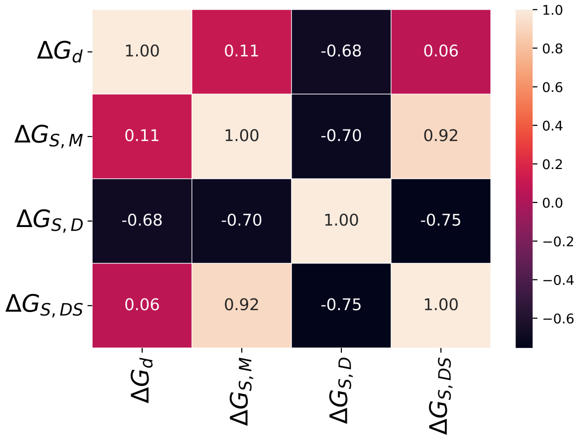

#### Figure S19. Fit of the model to ES and all ESI4c datasets

Curves in the same colors represent for the datasets in the same plates. Dots are the observed response. X axes are concentrations (M). The theoretical response $y_n * (\boldsymbol{\theta}^{MAP})$ is represented by the dashed line, where $\boldsymbol{\theta}^{MAP}$ is the MAP estimate, the mean of the posterior prediction is solid line, and the 95% posterior predictive interval is the shaded region. For the first plot: 100 nM (brown), 50 nM (red), and 25 nM (black) of the enzyme. For other plots: enzyme 100 nM, substrate 1350 nM (blue); enzyme 100 nM, substrate 750 nM (orange); enzyme 50 nM, substrate 150 nM (purple); enzyme 100 nM, substrate 50 nM (green).

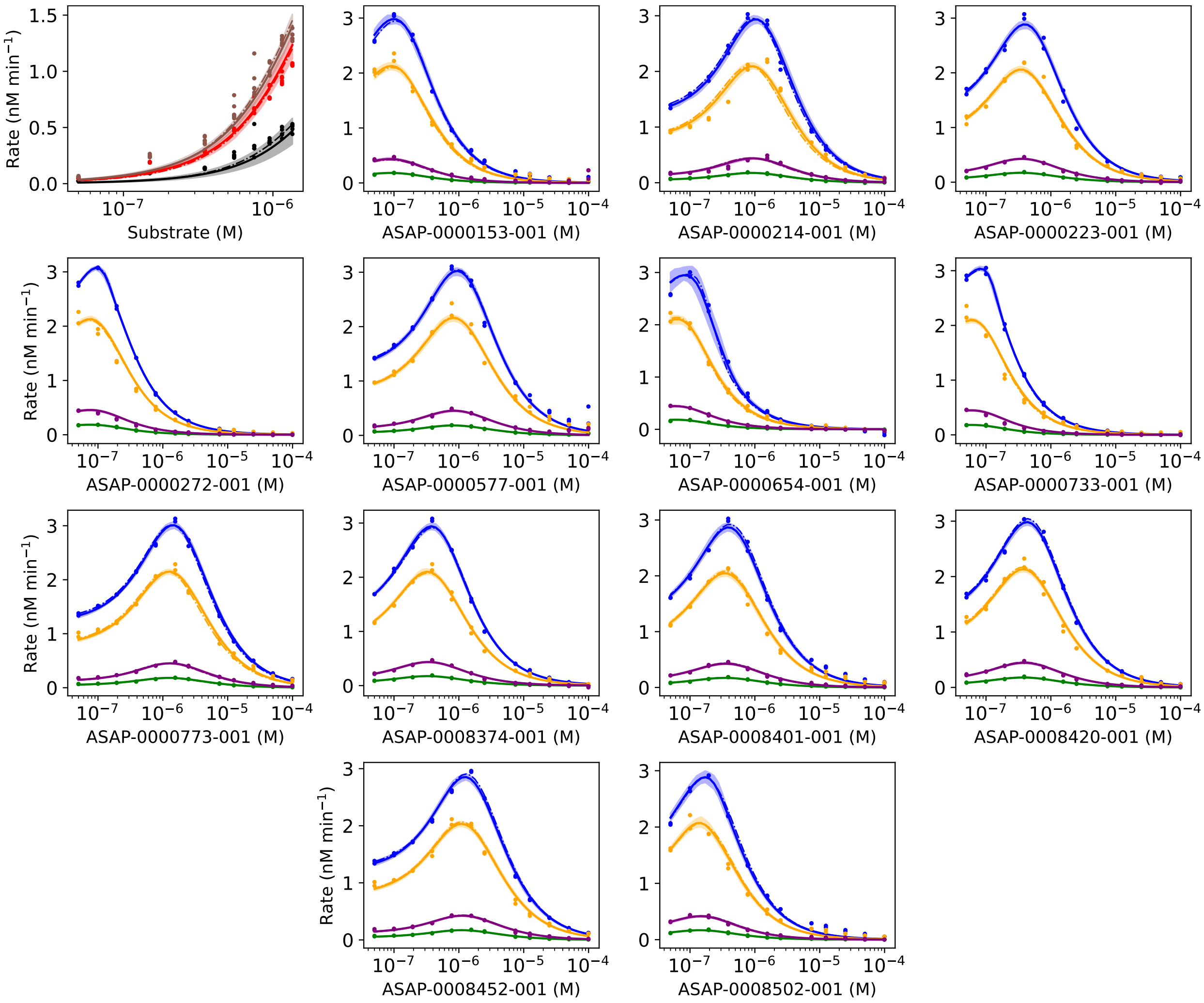

### Fitting of ES, all ESI4c, and 3 ESI1c datasets

#### Figure S20. 2D joint marginal distributions for pairs of some binding free energies of fitting ES, all ESI4c, and 3 ESI1c datasets

2D joint marginal probability densities were estimated based on 1,000 MCMC samples generated from the Bayesian posterior for all datasets.

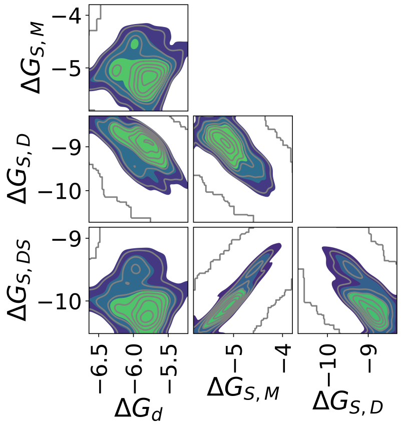

#### Figure S21. Heat map of the correlation matrix estimated from the Bayesian posterior for some binding free energies for fitting ES, all ESI4c, and 3 ESI1c datasets

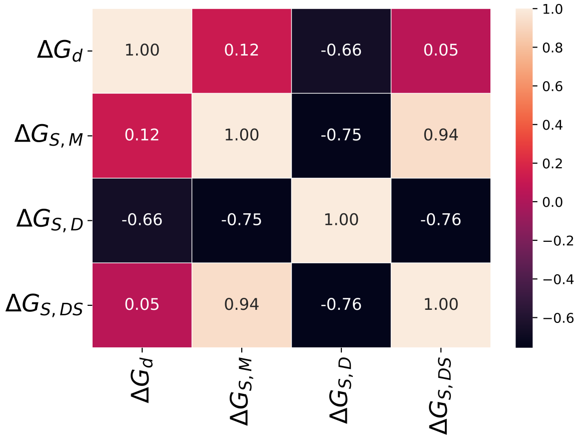

#### Figure S22. Ratios of rate constants of MERS-CoV MPro

Prior distributions are shown in black line, posterior of fitting datasets of ES + one ESI4c as green dashdot line, ES + all ESI4c as orange dashed line, ES + all ESI4c + three ESI1c as blue solid line. Red lines represent 95% HDI. The red triangle marks the median of the posterior.

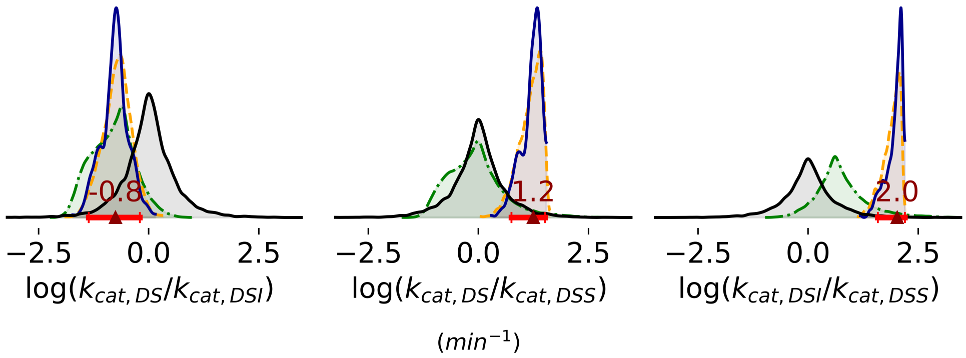

### Fitting 1 ESI1c dataset

#### Figure S23. Fit of the model to ESI1c datasets

Dots are the observed response. X axes are concentrations (M). The theoretical response $y_n * (\boldsymbol{\theta}^{MAP})$ is represented by the dashed line, where $\boldsymbol{\theta}^{MAP}$ is the MAP estimate, the mean of the posterior prediction is solid line, and the 95% posterior predictive interval is the shaded region.

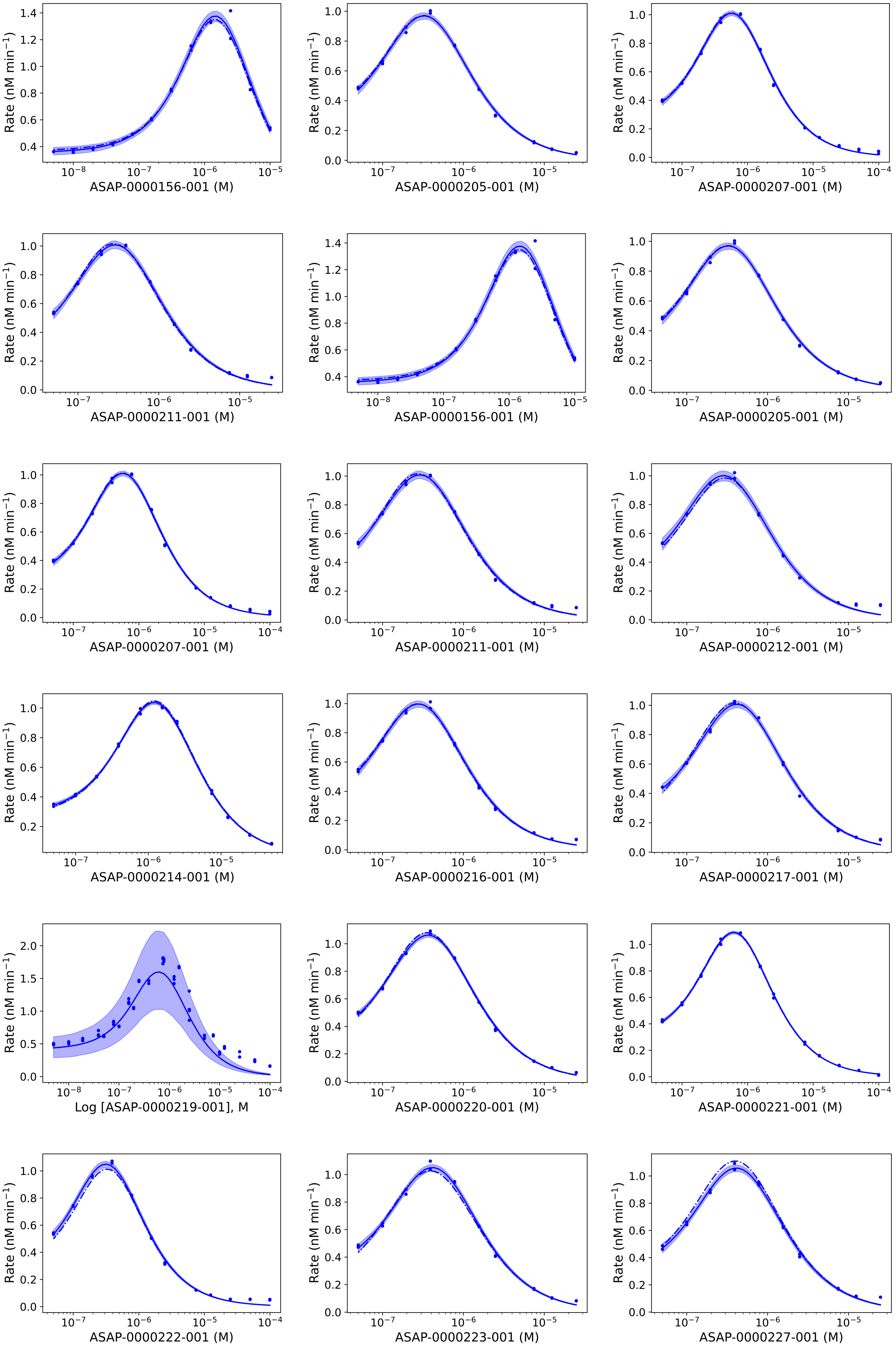
Rate (nM min$^{-1}$)
ASAP-0000156-001 (M)
ASAP-0000205-001 (M)
ASAP-0000207-001 (M)
ASAP-0000211-001 (M)
ASAP-0000156-001 (M)
ASAP-0000205-001 (M)
ASAP-0000207-001 (M)
ASAP-0000211-001 (M)
ASAP-0000212-001 (M)
ASAP-0000214-001 (M)
ASAP-0000216-001 (M)
ASAP-0000217-001 (M)
Log [ASAP-0000219-001], M
ASAP-0000220-001 (M)
ASAP-0000221-001 (M)
ASAP-0000222-001 (M)
ASAP-0000223-001 (M)
ASAP-0000227-001 (M)

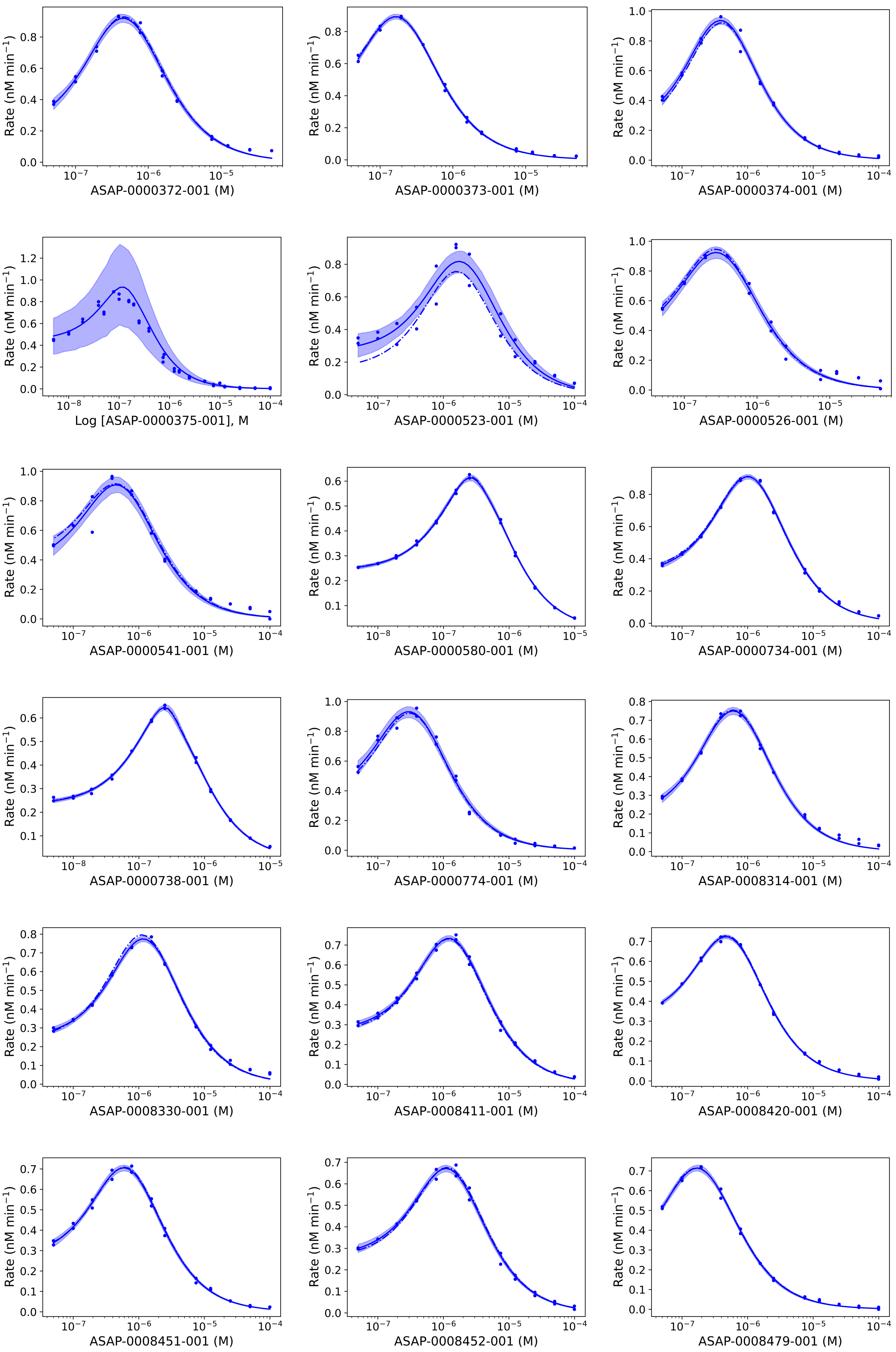
Rate (nM min$^{-1}$)
ASAP-0000372-001 (M)
ASAP-0000373-001 (M)
ASAP-0000374-001 (M)
Log [ASAP-0000375-001], M
ASAP-0000523-001 (M)
ASAP-0000526-001 (M)
ASAP-0000541-001 (M)
ASAP-0000580-001 (M)
ASAP-0000734-001 (M)
ASAP-0000738-001 (M)
ASAP-0000774-001 (M)
ASAP-0008314-001 (M)
ASAP-0008330-001 (M)
ASAP-0008411-001 (M)
ASAP-0008420-001 (M)
ASAP-0008451-001 (M)
ASAP-0008452-001 (M)
ASAP-0008479-001 (M)

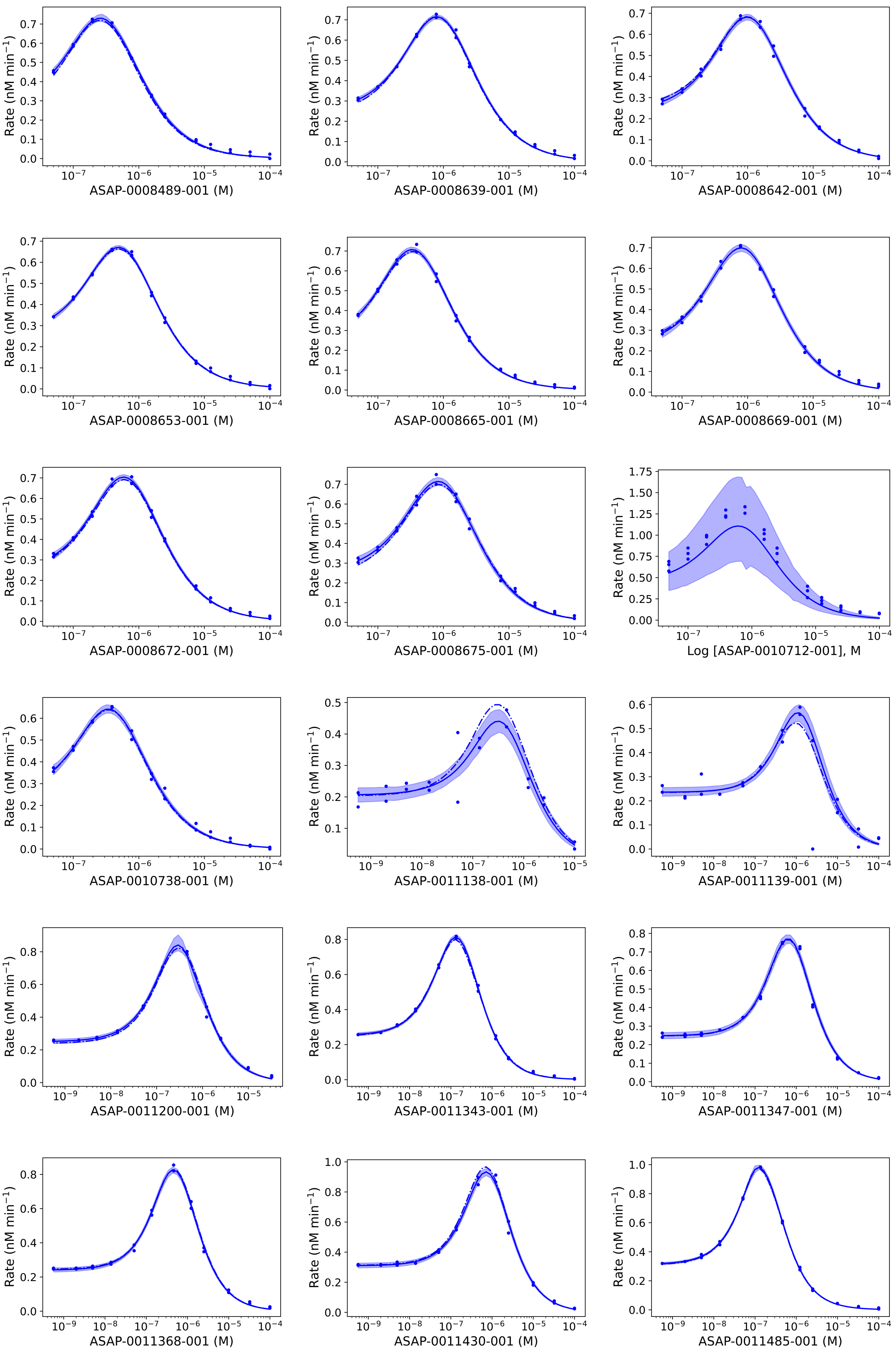
Rate (nM min$^{-1}$)
ASAP-0008489-001 (M)
ASAP-0008639-001 (M)
ASAP-0008642-001 (M)
ASAP-0008653-001 (M)
ASAP-0008665-001 (M)
ASAP-0008669-001 (M)
ASAP-0008672-001 (M)
ASAP-0008675-001 (M)
Log [ASAP-0010712-001], M
ASAP-0010738-001 (M)
ASAP-0011138-001 (M)
ASAP-0011139-001 (M)
ASAP-0011200-001 (M)
ASAP-0011343-001 (M)
ASAP-0011347-001 (M)
ASAP-0011368-001 (M)
ASAP-0011430-001 (M)
ASAP-0011485-001 (M)

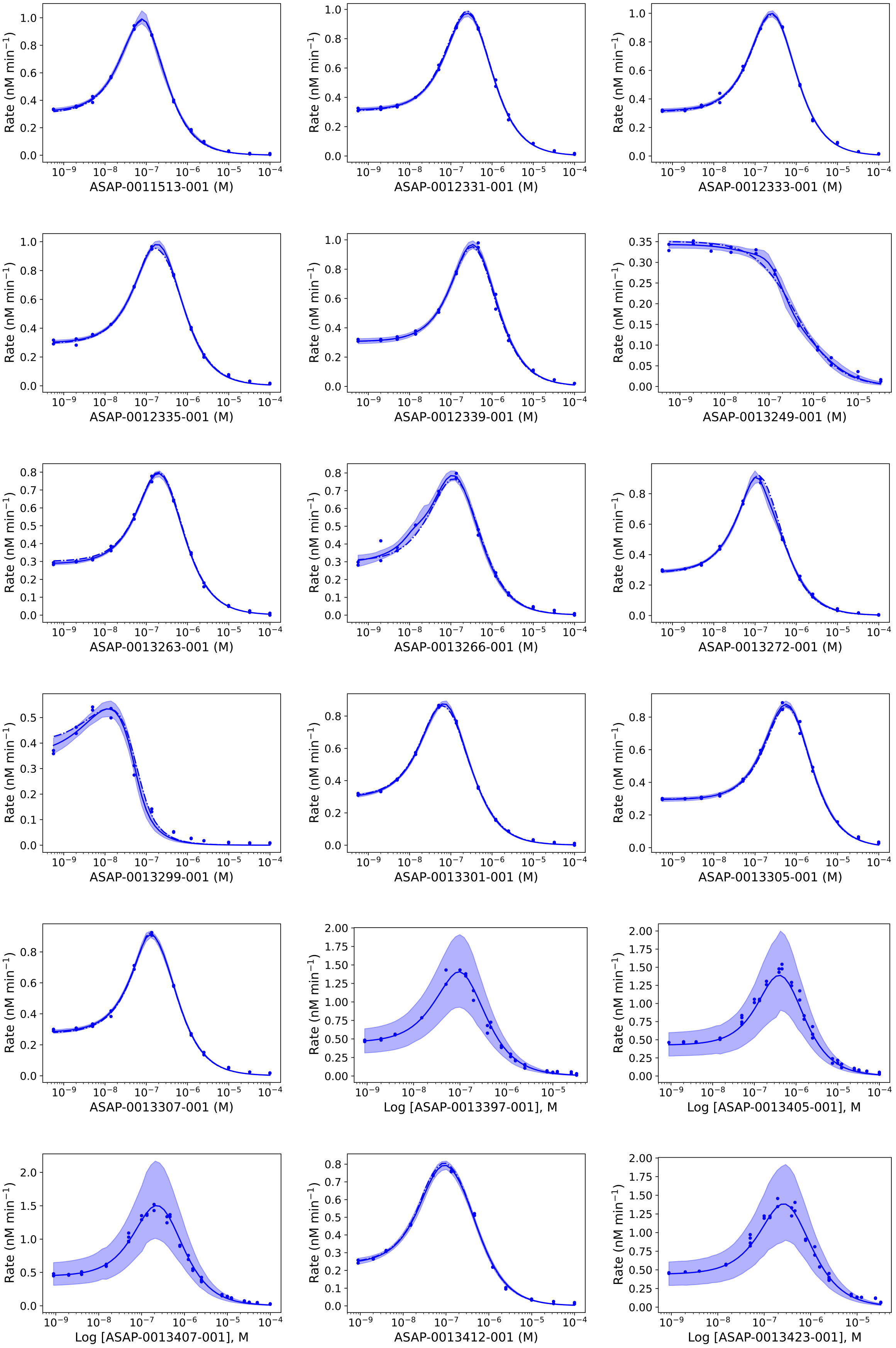
Rate (nM min$^{-1}$)
ASAP-0011513-001 (M)
ASAP-0012331-001 (M)
ASAP-0012333-001 (M)
ASAP-0012335-001 (M)
ASAP-0012339-001 (M)
ASAP-0013249-001 (M)
ASAP-0013263-001 (M)
ASAP-0013266-001 (M)
ASAP-0013272-001 (M)
ASAP-0013299-001 (M)
ASAP-0013301-001 (M)
ASAP-0013305-001 (M)
ASAP-0013307-001 (M)
Log [ASAP-0013397-001], M
Log [ASAP-0013405-001], M
Log [ASAP-0013407-001], M
ASAP-0013412-001 (M)
Log [ASAP-0013423-001], M

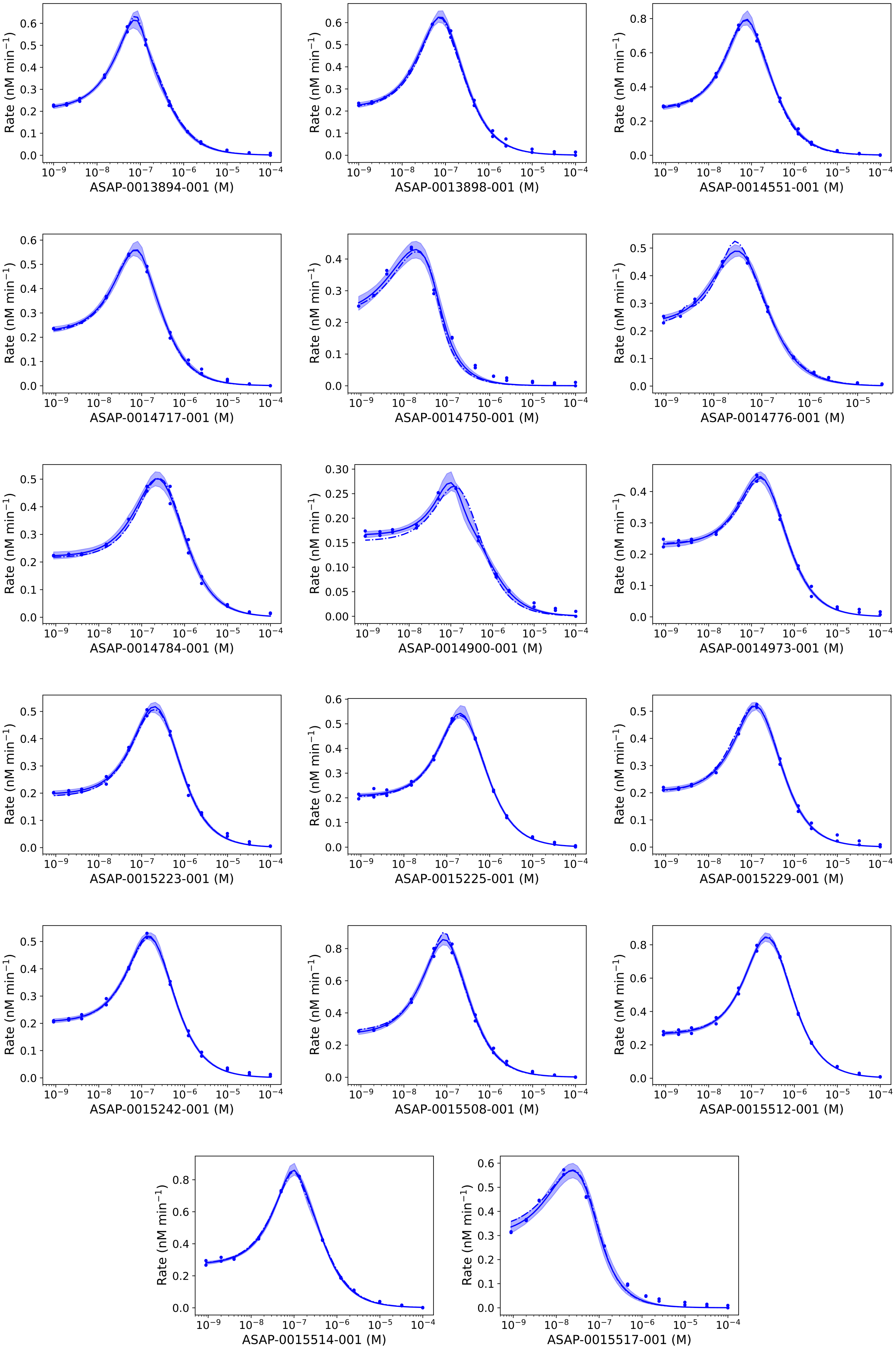
Rate (nM min$^{-1}$)
ASAP-0013894-001 (M)
ASAP-0013898-001 (M)
ASAP-0014551-001 (M)
ASAP-0014717-001 (M)
ASAP-0014750-001 (M)
ASAP-0014776-001 (M)
ASAP-0014784-001 (M)
ASAP-0014900-001 (M)
ASAP-0014973-001 (M)
ASAP-0015223-001 (M)
ASAP-0015225-001 (M)
ASAP-0015229-001 (M)
ASAP-0015242-001 (M)
ASAP-0015508-001 (M)
ASAP-0015512-001 (M)
ASAP-0015514-001 (M)
ASAP-0015517-001 (M)

#### Correlation Analysis

**Figure S24. Correlogram of inhibition, control, dimer pIC90 and cellular pEC90**

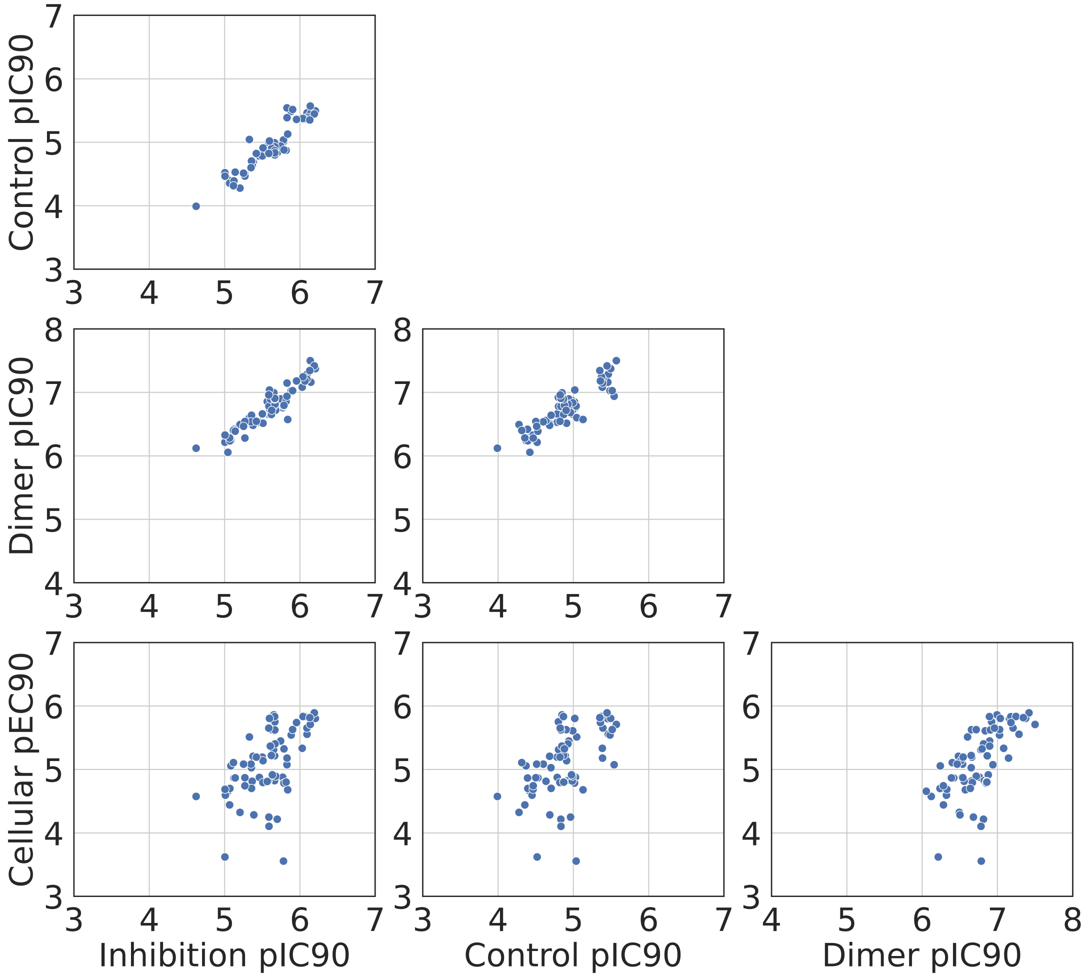

### Appendix C. MPro is primarily monomeric

Under the biochemical assay conditions, MERS MPro is primarily a monomer (Figure S25). Higher counts at larger mass than the monomer peak suggest that some of the dimeric form may be present in all the solutions. However, DiscoverMP only identifies a dimeric peak upon addition of a reversible covalent inhibitor, providing evidence of ligand-induced dimerization.

#### Figure S25. Mass photometry of 20 nM MPro

in biochemical assay conditions with (a) itself, (b) 7.5 $\mu$M substrate, and (c) 10 $\mu$M ASAP-0000214.

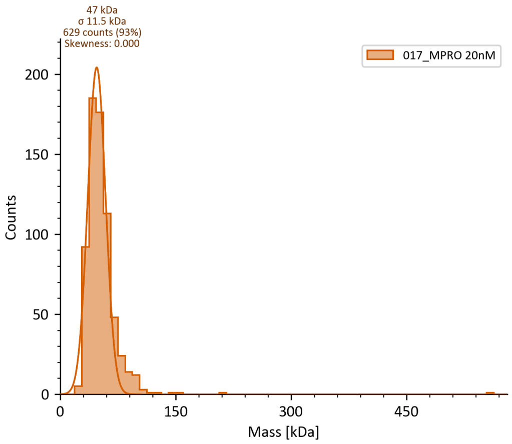

(a) MPro

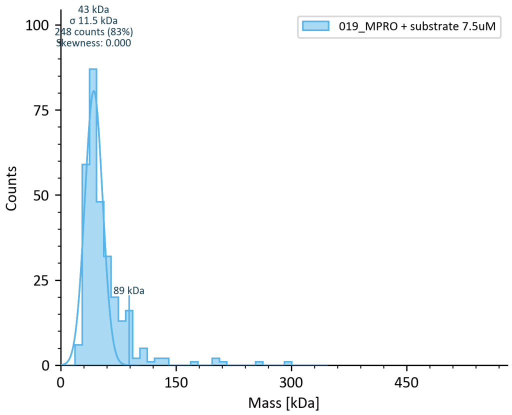

(b) MPro with substrate

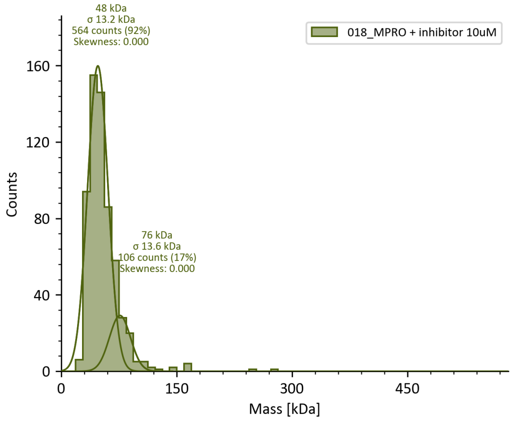

(c) MPro with 10 $\mu$M ASAP-0000214

#### Table S1. Correlation matrix of biochemical pIC90 and cellular pEC90 by Pearson R, Spearman $\rho$, and Kendall $\tau$

| Pearson R | Inhibition pIC90 | Control pIC90 | Dimer pIC90 |
|---|---|---|---|
| Control $pIC90$ | 0.954 ± 1.500E-2 | | |
| Dimer $pIC90$ | 0.873 ± 7.405E-2 | 0.858 ± 6.712E-2 | |
| Cellular $pEC90$ | 0.680 ± 7.374E-2 | 0.670 ± 7.343E-2 | 0.698 ± 6.266E-2 |

| Spearman $\rho$ | Inhibition pIC90 | Control pIC90 | Dimer pIC90 |
|---|---|---|---|
| Control $pIC90$ | 0.910 ± 2.578E-2 | | |
| Dimer $pIC90$ | 0.896 ± 5.474E-2 | 0.861 ± 5.055E-2 | |
| Cellular $pEC90$ | 0.685 ± 6.430E-2 | 0.639 ± 7.507E-2 | 0.748 ± 4.883E-2 |

| Kendall $\tau$ | Inhibition pIC90 | Control pIC90 | Dimer pIC90 |
|---|---|---|---|
| Control $pIC90$ | 0.754 ± 4.001E-2 | | |
| Dimer $pIC90$ | 0.776 ± 5.488E-2 | 0.698 ± 5.590E-2 | |
| Cellular $pEC90$ | 0.505 ± 5.642E-2 | 0.466 ± 6.426E-2 | 0.554 ± 4.683E-2 |

#### Table S2. Correlation matrix of biochemical pIC90 and cellular pEC90 by RMSD and aRMSD

| RMSD | Inhibition pIC90 | Control pIC90 | Dimer pIC90 |
|---|---|---|---|
| Control $pIC90$ | 0.684 ± 1.565E-2 | | |
| Dimer $pIC90$ | 0.613 ± 5.828E-2 | 0.296 ± 8.985E-2 | |
| Cellular $pEC90$ | 0.623 ± 6.124E-2 | 0.477 ± 3.895E-2 | 0.447 ± 5.074E-2 |

| aRMSD | Inhibition pIC90 | Control pIC90 | Dimer pIC90 |
|---|---|---|---|
| Control $pIC90$ | 0.132 ± 1.499E-2 | | |
| Dimer $pIC90$ | 0.250 ± 1.011E-1 | 0.264 ± 8.720E-2 | |
| Cellular $pEC90$ | 0.416 ± 5.399E-2 | 0.425 ± 5.247E-2 | 0.429 ± 5.921E-2 |

#### Table S3. p-value in comparison of correlation for $pIC50$s

| Pearson R | Cellular vs. Inhibition | Cellular vs. Control |
|---|---|---|
| Cellular vs. Control | 0.191 | |
| Cellular vs. Dimer | 0.181 | 0.242 |

| Spearman $rho$ | Cellular vs. Inhibition | Cellular vs. Control |
|---|---|---|
| Cellular vs. Control | 0.121 | |
| Cellular vs. Dimer | 0.018 | 0.004 |

| Kendall $\tau$ | Cellular vs. Inhibition | Cellular vs. Control |
|---|---|---|
| Cellular vs. Control | 0.171 | |
| Cellular vs. Dimer | 0.039 | 0.0191 |

| RMSD | Cellular vs. Inhibition | Cellular vs. Control |
|---|---|---|
| Cellular vs. Control | 8.919E-22 | |
| Cellular vs. Dimer | 1.694E-10 | 0.250 |

| aRMSD | Cellular vs. Inhibition | Cellular vs. Control |
|---|---|---|
| Cellular vs. Control | 0.037 | |
| Cellular vs. Dimer | 2.174E-04 | 0.002 |

#### Table S4. p-value in comparison of correlation for $pIC90$s

| Pearson R | Cellular vs. Inhibition | Cellular vs. Control |
|---|---|---|
| Cellular vs. Control | 0.184 | |
| Cellular vs. Dimer | 0.128 | 0.077 |

| Spearman $rho$ | Cellular vs. Inhibition | Cellular vs. Control |
|---|---|---|
| Cellular vs. Control | 0.026 | |
| Cellular vs. Dimer | 0.002 | 6.868E-06 |

| Kendall $\tau$ | Cellular vs. Inhibition | Cellular vs. Control |
|---|---|---|
| Cellular vs. Control | 0.027 | |
| Cellular vs. Dimer | 0.005 | 3.523E-05 |

| RMSD | Cellular vs. Inhibition | Cellular vs. Control |
|---|---|---|
| Cellular vs. Control | 1.959E-11 | |
| Cellular vs. Dimer | 1.204E-12 | 0.024 |

| aRMSD | Cellular vs. Inhibition | Cellular vs. Control |
|---|---|---|
| Cellular vs. Control | 0.168 | |
| Cellular vs. Dimer | 0.142 | 0.215 |

### Data

`SI.xlsx`

includes enzyme velocities for the datasets ES, ESI4c, and ESI1c.
